# Induction of viral mimicry upon loss of DHX9 and ADAR1 in breast cancer cells

**DOI:** 10.1101/2023.02.27.530307

**Authors:** Kyle A. Cottrell, Sua Ryu, Luisangely Soto Torres, Angela M. Schab, Jason D. Weber

## Abstract

Detection of viral double-stranded RNA (dsRNA) is an important component of innate immunity. However, many endogenous RNAs containing double-stranded regions can be misrecognized and activate innate immunity. The interferon inducible ADAR1-p150 suppresses dsRNA sensing, an essential function for ADAR1 in many cancers, including breast. Although ADAR1-p150 has been well established in this role, the functions of the constitutively expressed ADAR1-p110 isoform are less understood. We used proximity labeling to identify putative ADAR1-p110 interacting proteins in breast cancer cell lines. Of the proteins identified, the RNA helicase DHX9 was of particular interest. Knockdown of DHX9 in ADAR1-dependent cell lines caused cell death and activation of the dsRNA sensor PKR. In ADAR1-independent cell lines, combined knockdown of DHX9 and ADAR1, but neither alone, caused activation of multiple dsRNA sensing pathways leading to a viral mimicry phenotype. Together, these results reveal an important role for DHX9 in suppressing dsRNA sensing by multiple pathways.

## Introduction

RNA editing enhances protein diversity and modulates multiple aspects of RNA metabolism^1–3^. A-to-I editing is carried out by adenosine deaminase acting on RNA 1 (ADAR1), an RNA editase that binds double-stranded RNA (dsRNA) and converts adenosines to inosines^4,5^. The main domains of ADAR1 include the Z-DNA binding domains (ZBD), the double-stranded RNA (dsRNA) binding domains (dsRBD), and the deaminase domain^6,7^. There are two isoforms, p110 and p150, produced by alternative transcriptional start sites^8^. They share the same deaminase domain, dsRBDs, and a ZBD, but exhibit a distinct subcellular localization^9,10^. ADAR1-p150 is predominantly cytoplasmic, whereas ADAR1-p110 is nuclear-localized^9,11^.

An important role for ADAR1 is to suppress dsRNA sensing^12–15^. Many endogenously encoded RNAs can form large double-stranded regions, often through base-pairing between inverted Alu elements^16^. ADAR1 edits the majority of human genes, with most editing occurring within inverted Alu repeats^3,4^. By binding, editing, and thus altering the structure of dsRNA, ADAR1 suppresses the detection of dsRNA by various cytoplasmic sensors such as MDA5, RIG-I, and PKR^12,13,15,17,18^. These RNA sensors are part of innate immunity against viral infections^16,19–21^. Thus, ADAR1 prevents the activation of innate immunity pathways by endogenous immunogenic RNAs^16,20,22^. Some mutations of ADAR1 in humans cause inappropriate dsRNA sensing and activation of the type I IFN pathway which manifests as the interferonopathy Aicardi–Goutières syndrome^23^.

The cell-intrinsic antiviral response against foreign dsRNA – or misrecognized endogenous dsRNA – involves multiple pathways: (1) Recognition of dsRNA by MDA5 or RIG-I results in the activation of type I interferon (IFN-I) signaling^16,20,22^. (2) Activation of the dsRNA-binding kinase PKR triggers translational shutdown by phosphorylation of the translation initiation factor eIF2α^12^. (3) Detection of dsRNA by OAS proteins activates RNase L, which carries out non-specific cleavage of RNA and triggers cell death^24^. There is significant crosstalk between the three pathways, and the aforementioned dsRNA sensors are transcriptionally controlled by IFN-I, known as interferon-stimulated genes (ISGs)^25,26^. The ADAR1-p150 isoform is itself an ISG and is the isoform responsible for suppressing the activation of dsRNA sensors^8,10,14,27^. ADAR1-p110, however, is constitutively expressed, though its functions are less established^28,29^.

Because IFN-I signaling is often cytotoxic and antiproliferative, ADAR1’s ability to suppress IFN-I signaling was shown to exert pro-tumor effects^22,30,31^. As such, ADAR1 has been proposed as a potential therapeutic target for various cancers, including breast cancer^30,32–34^. ADAR1 mRNA expression is elevated in breast cancer and is correlated with a poor prognosis^30,35^. Furthermore, ADAR1 is essential for a subset of breast cancer cells overrepresented by triple-negative breast cancer (TNBC)^35^. Known as ‘ADAR1-dependency,’ depletion of ADAR1-p150 leads to IFN-I signaling and global translational repression in cells that are sensitive to depletion of ADAR1^34,35^. What makes cancer cells sensitive or refractory to ADAR1 loss is not yet determined^35^.

ADAR1 interacts with numerous RNA binding proteins including RNA helicases, transcription machinery, and DNA repair proteins^29,36–39^. The influence of ADAR1-interacting proteins on A-to-I editing has previously been reported^37,40–43^. In this study, we evaluated components of the ADAR1 interactome in breast cancer cells and identified the RNA helicase DHX9 as a redundant suppressor of immunogenic dsRNA in ADAR1-independent breast cancer cells. We demonstrate that co-depletion of ADAR1 and DHX9 is sufficient to trigger a viral mimicry phenotype in ADAR1-independent cells.

## Results

### Proximity Labeling by APEX2 Reveals ADAR1 Interacting Proteins

To better understand the role of ADAR1-p110 in breast cancer, we turned to a proximity labeling approach using APEX2 to identify putative ADAR1-p110-interacting proteins. Proximity labeling by APEX2 allows for the identification of proteins within 20 nm of an APEX2 fusion protein via biotin-mediated pulldown^44^. An APEX2-ADAR1-p110 fusion construct or APEX2 alone was expressed in MDA-MB-231, MCF-7, and SK-BR-3 (Figure 1a). Following proximity labeling, biotinylated proteins were purified by streptavidin pulldown and subsequently identified by mass spectrometry (Fig 1b-c, Figure S1a, Supplementary Table 2). In total, we identified over one hundred enriched proteins across the three cell lines, with many identified in all three lines (Figure 1d). Over-representation analysis of gene ontology (GO) terms revealed that many of the proteins identified by proximity labeling have roles in multiple aspects of RNA metabolism and localize to the nucleus and nucleolus (Figure 1e, Supplementary Table 3). This finding was supported by a comparison to proteins previously observed to localize to the nucleus and nucleolus within MCF-7 in the SubCellBarcode dataset^45^ (Figure 1f and 1g). These findings are largely consistent with the localization of ADAR1 within the cell lines studied. Immunofluorescence for ADAR1, which largely reflects the localization of ADAR1-p110 as the predominant isoform, showed that ADAR1 was localized in both the nucleus and nucleolus (Figure 1h).

**Figure 1:**
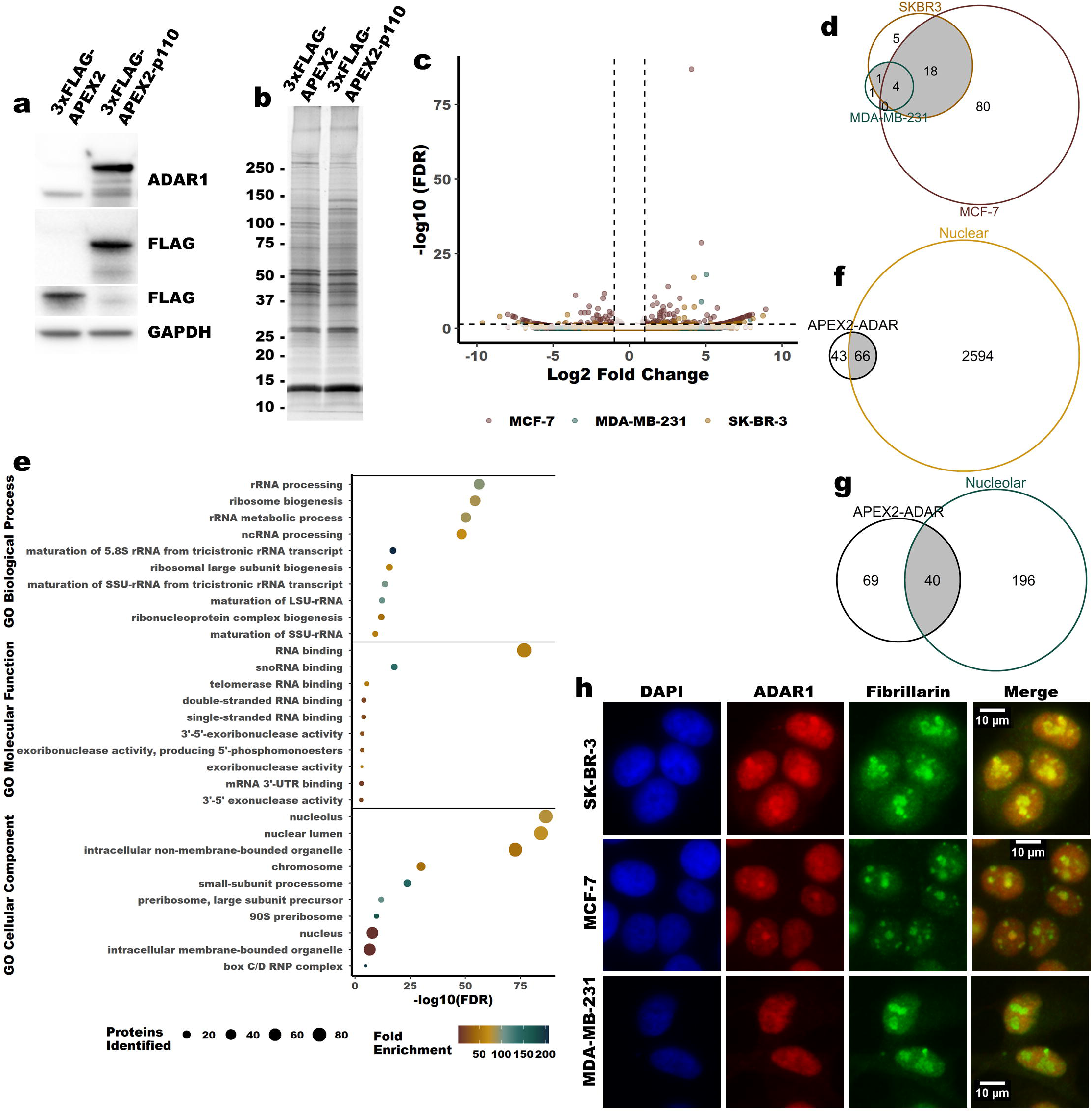
Identification of putative ADAR1 interacting proteins by APEX2 proximity labeling. **a** Representative immunoblot showing expression of the constructs used for proximity labeling in MCF-7, MDA-MB-231 and SK-BR-3, MDA-MB-231 is shown. **b** Representative fluorescently stained gel image showing proteins purified by streptavidin-biotin pulldown following proximity labeling in MCF-7. **c** Volcano plot summarizing the proteins identified by mass spectrometry following streptavidin pulldown subsequent to proximity labeling. Differential abundance of all proteins in each cell line can be found in Supplementary Table 2. **d** Venn diagram showing overlap between enriched proteins from all three cell lines. The cut-off for enriched proteins was an FDR adjusted p-value of less than 0.05 and a Log2 fold change of greater than 0.5. **e** GO terms found to be overrepresented in the list of the enriched proteins. Only the top ten GO terms, by FDR, are shown for each category. All other significant GO terms are available in the Supplementary Table 3-4. **f** and **g** Venn diagrams showing overlap between the enriched proteins identified and those proteins found previously to localize to the nucleus or nucleolus in MCF-7^45^. **h** Representative indirect immunofluorescence micrographs showing localization of ADAR1, fibrillarin (a nucleolar marker) and DAPI.

### Validation of Protein Interactions Identified by Proximity Labeling

Although proximity labeling by APEX2 is a powerful technique for identifying putative protein-protein interactions, it does not distinguish between interactions and close associations^44^. To validate the proximity labeling findings and provide supporting evidence for direct protein-protein interactions, we performed co-immunoprecipitation followed by immunoblotting for five proteins identified by proximity labeling. ADAR1 could be immunoprecipitated by antibodies against the helicases DHX9, DDX17, and DDX54 (Figure 2a-g). ADAR1 could also be immunoprecipitated with antibodies against XRN2 and PARP (Figure S1b-c). To assess whether these potential interactions depended on RNA, we treated the lysates with RNase A to degrade RNA prior to immunoprecipitation (Figure S1d). Immunoprecipitation of ADAR1 by antibodies against DHX9, DDX17, and DDX54 was possible even in the presence of RNase A and improved in some cases (Figure 2a-g). For DDX17 and DDX54, RNase A treatment did not change the co-immunoprecipitation results. However, DHX9 immunoprecipitated with both isoforms of ADAR1 in the absence of RNase A, but only with ADAR1-p110 when treated with RNase A. These findings suggest that DHX9, DDX17, and DDX54 directly interact with ADAR1-p110, and DHX9 interacts with ADAR1-p150 through an RNA bridge/scaffold. The co-immunoprecipitation findings are also consistent with the localization of the proteins studied. Much like ADAR1, the helicases DHX9, DDX17, and DDX54 localized to the nucleus or nucleolus (Figure 2h-j). Interestingly, in the cell lines used in this study, ADAR1-p150 is localized to the nucleus and cytosol (Figure S1e), consistent with previous reports of ADAR1-p150 shuttling between these two compartments^46,47^. Together, these findings validate the results of the proximity labeling described in Figure 1 and provide evidence for direct interactions between ADAR1 and multiple helicases.

**Figure 2:**
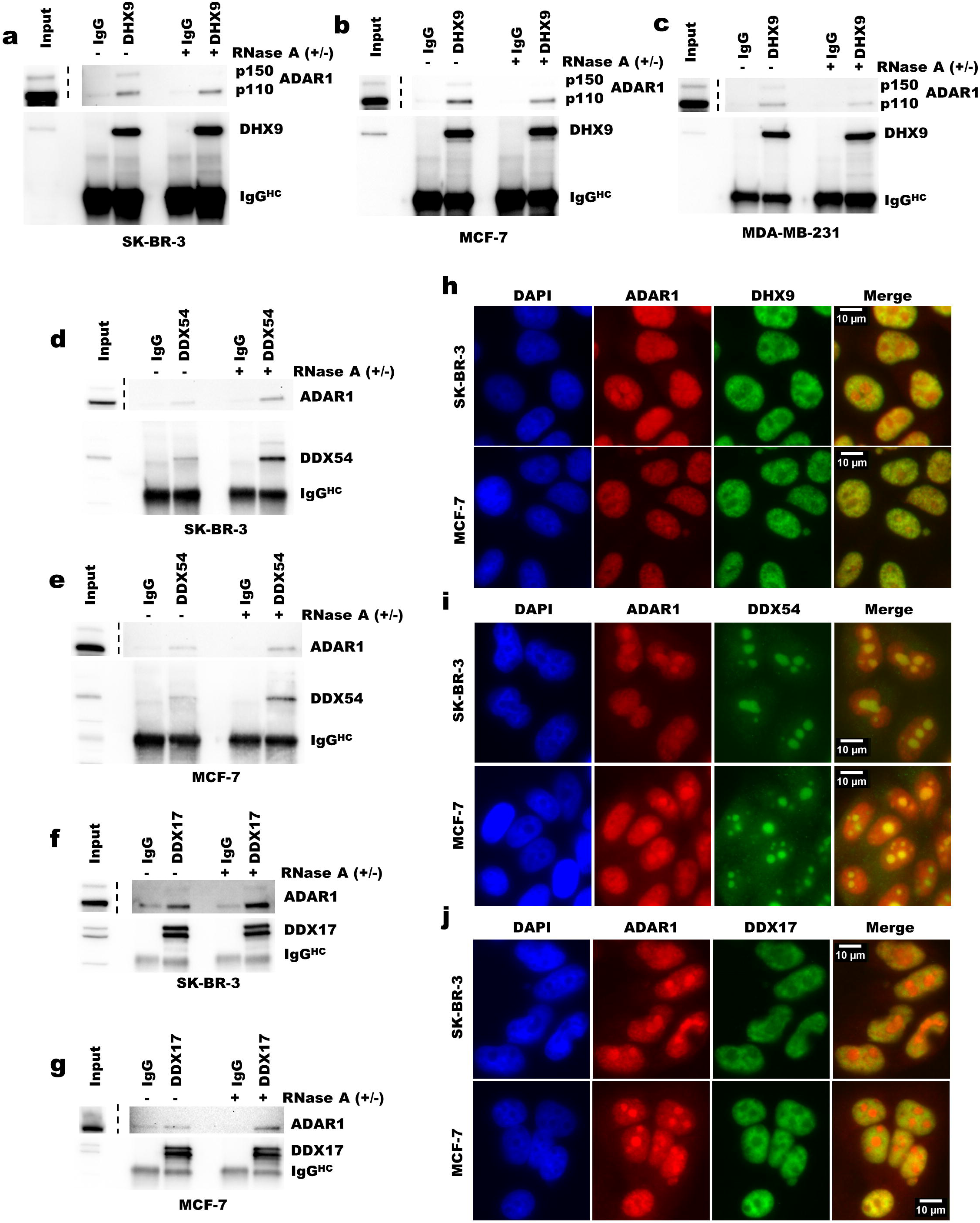
Validation of putative protein-protein interactions identified by proximity labeling Immunoprecipitation of DHX9. **a-c**, DDX54 **d**-**e** or DDX17 **f**-**g** followed by immunoblot for ADAR in breast cancer cell lines. Immunoblot of immunoprecipitation eluates and inputs from SK-BR-3 **a**, **d** and **f**, MCF-7 **b**, **e** and **g**, and MDA-MB-231 **c**. Input represents 5% of the lysate used for immunoprecipitation. The IgG lanes represent immunoprecipitation eluates from pulldown with anti-rabbit IgG antibody. The lanes labeled DHX9, DDX54 and DDX17 indicate the eluates from immunoprecipitation with antibodies against those proteins respectively. The IgG^HC^ label indicates the band corresponding to the IgG heavy chain from the antibody used for immunoprecipitation. Uncropped immunoblots for panels **a**-**g** can be found in Source Data Figures. Immunofluorescence for ADAR1 and DHX9 **h**, DDX54 **i**, or DDX17 **j** in SKBR3 or MCF-7.

### DHX9 is overexpressed in breast cancer

Of the identified helicases, DHX9 was of particular interest for several reasons. First, DHX9 is the only helicase in humans that has a dsRBD. The dsRBD that is present in DHX9 is of the same class present in ADAR1 and PKR (Figure 3a). Analysis of publicly available RNA-seq data for human cell lines and tumors revealed that DHX9 expression closely correlates with ADAR1 expression (Figure 3b-d, Figure S2a-c). The correlation is stronger between DHX9 and the transcript encoding ADAR1-p110, than between DHX9 and ADAR1-p150 (Figure 3c-d, Figure S2d-e). Furthermore, the expression of DHX9 and ADAR1 correlates better than any other helicase identified by proximity labeling (Figure 3b, Figure S2b). Consistent with this correlated expression, and much like ADAR1, DHX9 is highly expressed in breast cancer and is correlated with a poor prognosis^35^ (Figure 3c, Figure S2f-h).

**Figure 3:**
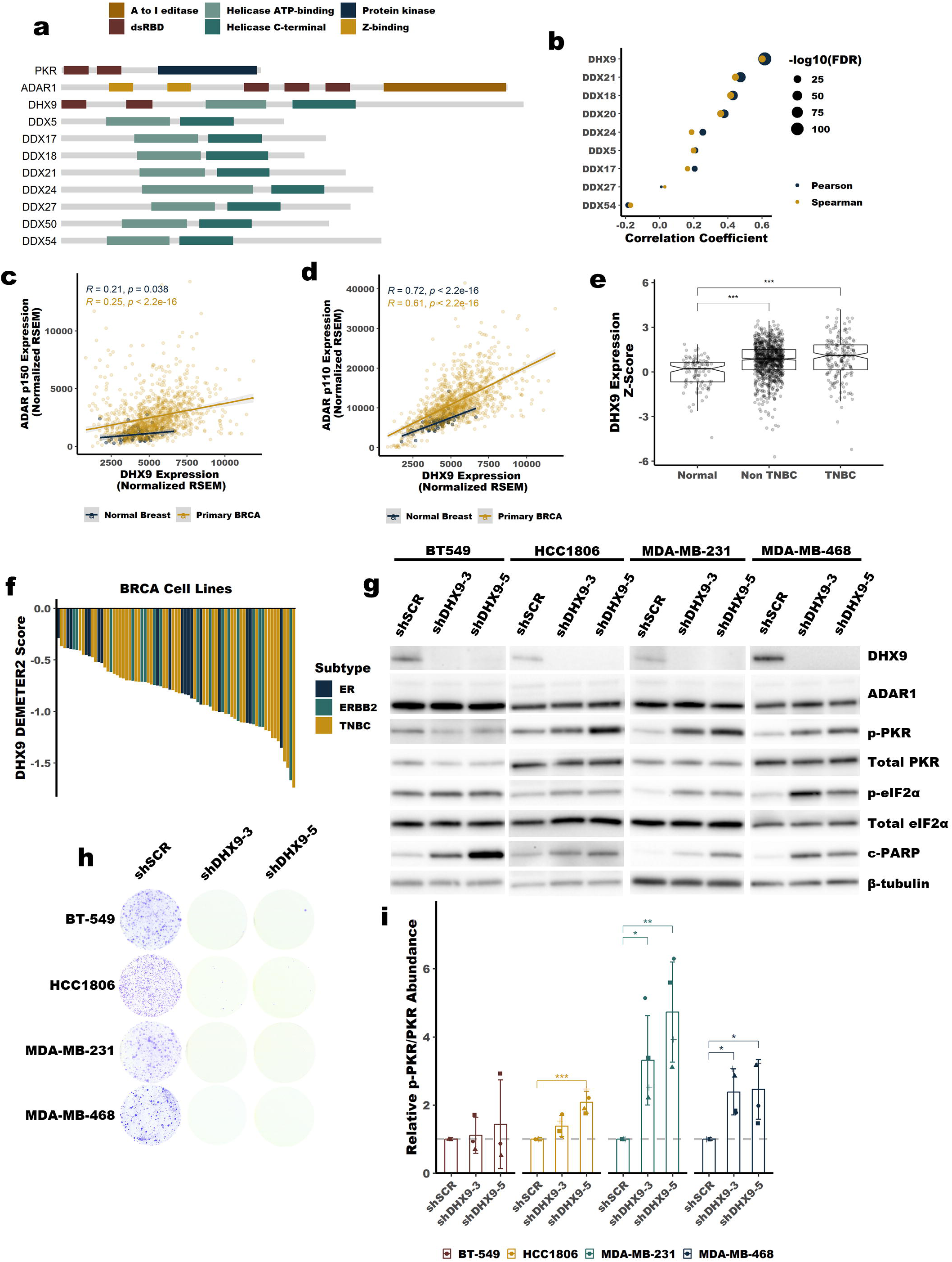
DHX9 is overexpressed in breast cancer and suppresses PKR activation. **a** Schematic showing the domain structure of PKR, ADAR1, DHX9 and other helicases identified by proximity labeling in Figure 1, dsRBD refers to the dsRNA Binding Domain. **b** Pearson and Spearman correlation coefficients for the correlation between ADAR1 expression at the RNA level and the expression of each indicated helicase at the RNA level, data from breast tumors within TCGA. Scatterplots showing the correlation between ADAR1-p110 **c**, or ADAR1-p150 **d**, and DHX9 expression in normal breast or breast tumors. **e** Expression of DHX9 at the RNA level in normal breast, non-TNBC or TNBC tumors. **f** Waterfall plot showing DHX9 dependency of breast cancer cell lines using data from DepMap, ER = estrogen receptor positive cell lines, ERRB2 = HER2 positive cell lines. **g** Representative immunoblot following knockdown of DHX9 with two different shRNAs in four TNBC cell lines. Immunoblots for other replicates and uncropped blots can be found in Source Data Figures. **h** Foci formation assay for the same cells used in **g** for immunoblot. **i** Quantification of PKR phosphorylation as determined by the immunoblot in **g**. Quantification of protein expression for other proteins of interest can be found in Figure S3a-e. Bars represent the average of at least three replicates, error bars are +/− SD. * p <0.05, ** p <0.01, *** p < 0.001. P-values determined by Dunnett’s test.

### DHX9 is essential in TNBC cell lines and suppresses PKR activation

The similarities between ADAR1 and DHX9 led us to further study the role of DHX9 in breast cancer. Analysis of publicly available data from DepMap (https://depmap.org/portal/download/custom/) revealed that DHX9 is commonly essential in breast cancer cell lines (Figure 3f). We validated this finding by knocking down DHX9 in four TNBC cell lines previously shown to be ADAR1-dependent^35^. In all four lines, knockdown of DHX9 caused cell death, likely through caspase-mediated apoptosis as indicated by cleaved PARP (Figure 3g and h). Given the presence of the common dsRBD in DHX9 and PKR, and the role of DHX9 in regulating the abundance of dsRNA^48^, we asked whether DHX9 could influence activation of PKR. To our surprise, we found that in three of the TNBC cell lines studied, knockdown of DHX9 caused activation of PKR (Figure 3g and 3i), this finding was confirmed for one cell line by CRISPR-Cas9 knockout of DHX9 (Figure S3i-k). Together these findings show that DHX9 is essential in breast cancer cell lines and that in some ADAR1-dependent cell lines, DHX9 suppresses PKR activation.

### DHX9 and ADAR1 redundantly suppress PKR activation

The experiments above were performed in ADAR1-dependent TNBC cell lines – cell lines that activate PKR following ADAR1 knockdown^35^. We were curious if DHX9 knockdown would also cause PKR activation in ADAR1-independent cell lines – cell lines that do not activate PKR following ADAR1 knockdown. Using shRNAs, we knocked down DHX9 in two ADAR1-independent breast cancer lines, MCF-7 and SK-BR-3 (Figure 4a-b and 4f-g, Figure S4a-b and 4g-h). Unlike in ADAR1-dependent cell lines, knockdown of DHX9 did not cause activation of PKR in ADAR1-independent cell lines (Figure 4a, 4d, 4f, 4i). Next, we asked whether combined knockdown of ADAR1 and DHX9 in these cell lines would lead to activation of PKR. As we had previously observed, knockdown of ADAR1 in SK-BR-3 and MCF-7 did not cause PKR activation^35^. However, combined knockdown of DHX9 and ADAR1 caused robust activation of PKR in both cell lines (Figure 4a, 4d, 4f, 4i). Consistent with PKR activation, we observed increased phosphorylation of the PKR substrate eIF2 [following combined knockdown of ADAR1 and DHX9 (Figure 4a and 4d, Figure S4c and S4i). Like in ADAR1-dependent TNBC cell lines, knockdown of DHX9 caused reduced proliferation of MCF-7 and SK-BR-3 (Figure 4c, 4d, 4h, 4j), likely through caspase-dependent apoptosis, as indicated by elevated cleaved PARP (Figure 4a, 4d, Figure S4e, S4k). Although PARP cleavage was increased upon combined knockdown of DHX9 and ADAR1, this did not statistically reduce the proliferation of the cells compared to single knockdown of ADAR1 and DHX9 as measured by foci formation. Together, these results reveal that ADAR1 and DHX9 redundantly suppress PKR activation in ADAR1-independent breast cancer cell lines.

**Figure 4:**
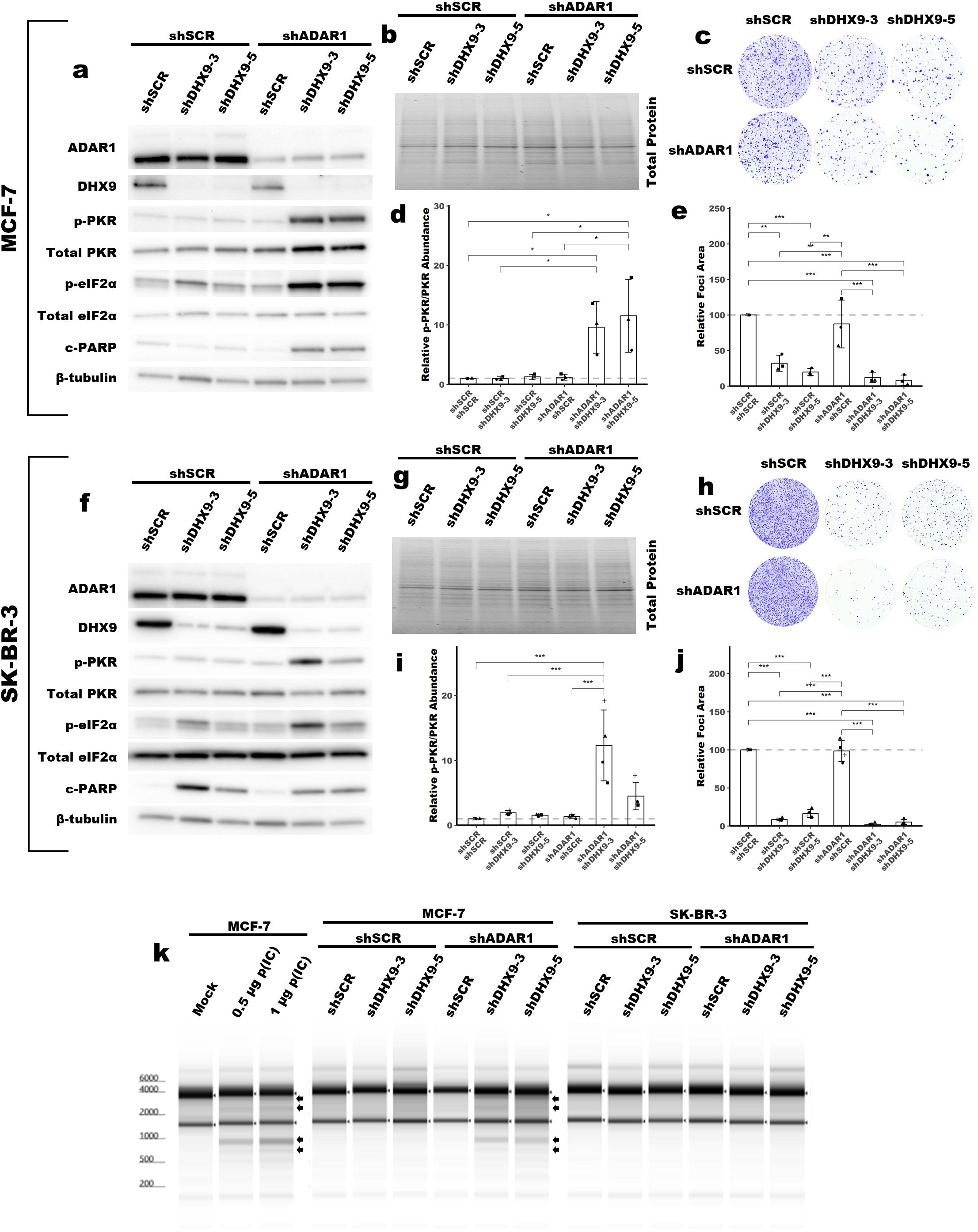
DHX9 and ADAR1 redundantly suppress dsRNA sensing in ADAR1-independent cell lines Representative immunoblot showing the phenotype of ADAR1 and/or DHX9 knockdown in MCF-7. **a**, or SK-BR-3 **f**. Immunoblots for other replicates and uncropped blots can be found in Source Data Figures. Protein abundance from the immunoblot in **a** and **f** was normalized by total protein abundance by quantification of the Stain Free gel in **b** and **g** respectively. Fold change of PKR phosphorylation at Thr-446 in MCF-7 **d** or SK-BR-3 **i** as determined by the immunoblots in **a** or **f** respectively. Quantification of protein expression for other proteins of interest can be found in Figure S4a-l. Representative foci formation phenotype of ADAR1 and/or DHX9 knockdown in MCF-7 **c** or SK-BR-3 **h**, quantification of relative foci area is shown in **e** or **j** respectively. **k** Analysis of rRNA integrity upon knockdown of ADAR1 and/or DHX9 in MCF-7 or SK-BR-3. Additionally, panel **k** shows the effect of poly(I:C) (p(I:C)) transfection on rRNA integrity in MCF-7. Arrows indicate canonical RNase L cleavage products^50^. Bars represent the average of at least three replicates, error bars are +/− SD. * p <0.05, ** p <0.01, *** p < 0.001. P-values determined by one-way ANOVA with post-hoc Tukey. Comparisons between the two different shRNAs targeting DHX9 (shDHX9-3 and shDHX9-5) were not included for clarity.

### DHX9 and ADAR1 redundantly suppress RNase L activation

Next, we wanted to evaluate the activation of other dsRNA sensing pathways in ADAR1-independent cell lines after the combined knockdown of ADAR1 and DHX9. To assess whether the IFN-I pathway, or other pathways, is activated after combined knockdown of ADAR1 and DHX9 we turned to analysis of differential gene expression by RNA-seq. In the process of preparing RNA for sequencing, we were surprised to find specific degradation of rRNA in MCF-7 cells following combined knockdown of ADAR1 and DHX9 (Figure 4k). Single knockdown of either DHX9 or ADAR1 did not cause rRNA degradation. The observed rRNA degradation in the combined knockdown cells is consistent with the degradation products caused by RNase L, as indicated in Figure 4k^49,50^. Transfection with poly(I:C), which activates the IFN-I pathway and RNase L^51^, created an identical band pattern to that of combined knockdown of DHX9 and ADAR1, indicating that the degradation of rRNA observed in these cells is likely caused by RNase L activity (Figure 4k). We performed the same experiment with the ADAR1-dependent TNBC cell lines described above. In these cells, we did not see activation of RNase L after the knockdown of DHX9 alone (Figure S3h).

### DHX9 and ADAR1 redundantly suppress multiple innate immunity pathways

RNA sequencing revealed that many more RNAs were differentially expressed after combined knockdown of DHX9 and ADAR1, compared to single knockdown of ADAR1 or DHX9 (Figure 5a, Figure S5a-f). Analysis of differential gene expression by gene set enrichment after combined knockdown of ADAR1 and DHX9 in MCF-7 revealed activation of multiple pathways involved in the innate response to viral infection and repression of several pathways involved in translation (Figure 5b and Supplemental Table 14). An enrichment map showed that the activated pathways associated with innate immunity formed one cluster and the depressed pathways formed a separate cluster (Figure S5g). Of the upregulated pathways, several were associated with activation of IFN signaling (Figure 5b, and Supplementary Table 14). Analysis of core ISG expression revealed significant upregulation of ISGs in MCF-7 after combined knockdown of ADAR1 and DHX9 (Figure 5c and Figure S5c). On the contrary, knockdown of DHX9 or ADAR1 alone did not increase ISG expression. Activation of RNase L described above also indicated that the type I IFN pathway was active in MCF-7 after double knockdown of DHX9 and ADAR1, because the activators of RNase L, the OAS proteins, are ISGs^24^. In fact, a GO term associated with OAS activity was upregulated in MCF-7 after combined knockdown of ADAR1 and DHX9 (Supplementary Table 14). Consistent with activation of PKR, we also observed increased expression of ATF4 targets (Figure 5d) and NF-KB targets (Figure 5e). Interestingly, we did not observe activation of any of these pathways or RNase L in SK-BR-3 following combined ADAR1 and DHX9 knockdown. It is not clear from these data what causes this difference between SK-BR-3 and MCF-7. SK-BR-3 expresses MDA-5, MAVS, and PKR to a similar level as MCF-7 (Supplemental Figure 6g). Furthermore, both cell lines respond to transfection of p(I:C) by activation of PKR and the type I IFN pathway as indicated by increased expression of PKR and MDA5 (Figure S6h).

**Figure 5:**
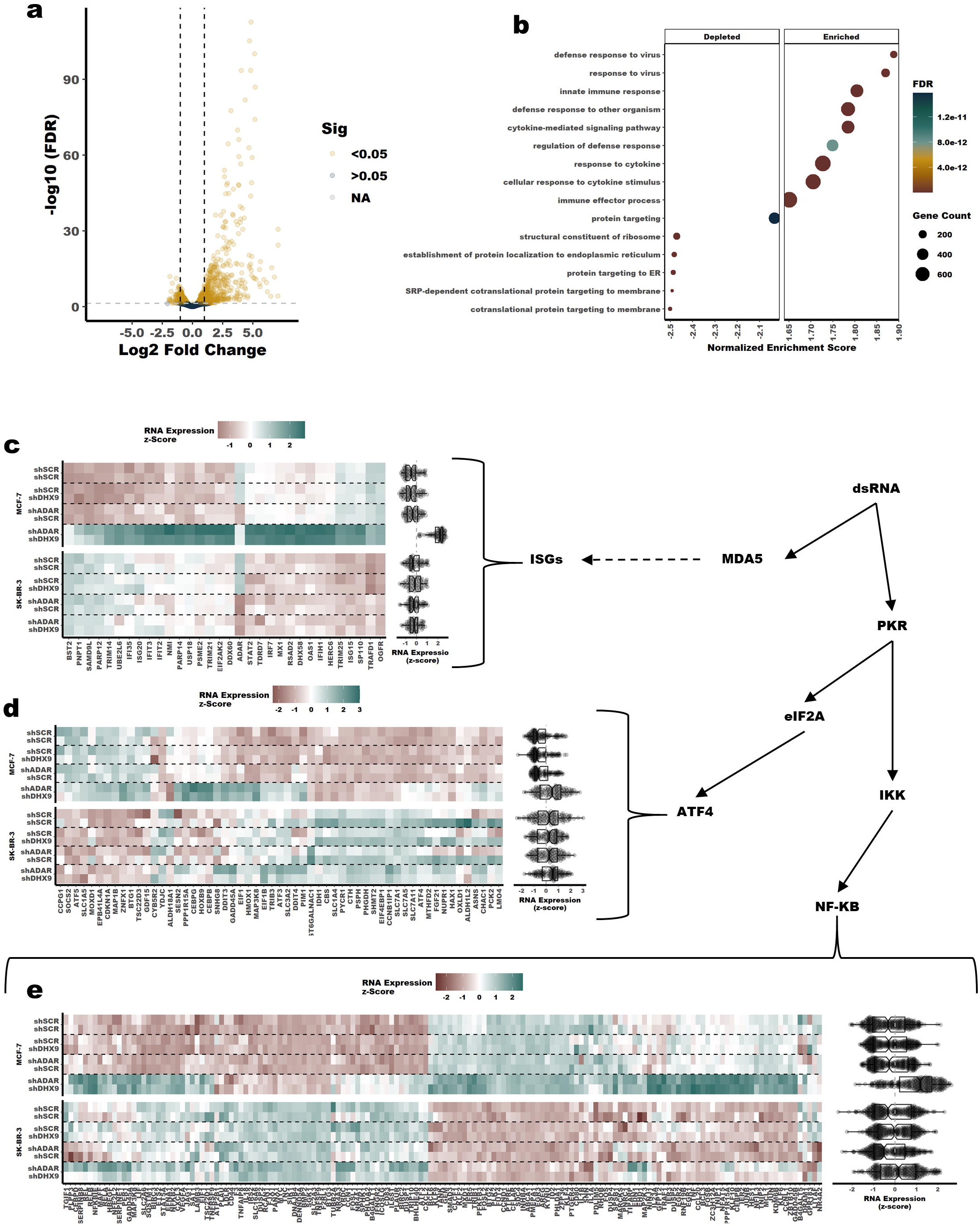
Induction of a viral mimicry phenotype upon knockdown of DHX9 and ADAR1 in MCF-7. **A** Volcano plot showing changes in RNA expression upon knockdown of DHX9 and ADAR1 in MCF-7, a volcano plot for SK-BR-3 is in Figure S5f. Fold-change of RNA expression shown in **a** was determined using an interaction term between ADAR1 and DHX9 knockdown, volcano plots for fold change of RNA expression for single knockdown of ADAR1 or DHX9 is in Figure S5a-b and S5d-e. **b** GO terms identified by gene set enrichment analysis of the RNA-seq data in **a**. **c**-**e** Heatmaps and summary box plots showing RNA expression changes in MCF-7 and SK-BR-3 upon knockdown of ADAR1 and/or DHX9. Panel **c** shows RNA expression for core ISGs^22,35^, panel **d** shows ATF4 targets and panel **e** shows NF-KB targets with ISGs removed.

The finding that combined knockdown of ADAR1 and DHX9 did not induce ISG expression in SK-BR-3 is consistent with our findings in TNBC cell lines. While knockdown of DHX9 alone caused activation of PKR in several TNBC cell lines, we observed no activation of the type I IFN pathway, as indicated by no change in ISG15 expression (Figure S3f-g). ISG15 was found to be highly upregulated at the RNA and protein level in MCF-7 after double knockdown of ADAR1 and DHX9 (Figure S4f, S4m, Supplementary Table 8). Like ISG expression overall, ISG15 expression in SK-BR-3 was not changed by knockdown of DHX9 and ADAR1 (Figure S4l, S4n).

Given the previously described role of DHX9 in the control of Alu containing RNAs^48^, we next sought to assess if increased expression of transposable elements, especially Alus, could explain the activation of PKR or the IFN pathway upon combined knockdown of DHX9 and ADAR1. Analysis of our RNA-seq data revealed that transposable element expression was generally unchanged upon either single knockdown of ADAR1 or DHX9, or combined knockdown of both proteins (Figure S6a-f).

### The dsRBDs of DHX9 are sufficient to suppress PKR activation in the absence of ADAR1

Having shown through knockdown studies that ADAR1 and DHX9 function redundantly to suppress dsRNA sensing, we next wanted to assess which functions of DHX9 and which isoforms of ADAR1 are important for this role. To determine which functions of DHX9 are sufficient to suppress PKR activation, we performed a rescue experiment with wild-type DHX9, a helicase deficient mutant DHX9, and a truncated DHX9 that possess the N-terminal dsRBDs fused to EGFP (Figure 6a, Figure S6a). Overexpression of wild-type DHX9 rescued the PKR activation phenotype caused by double knockdown of ADAR1 and DHX9, confirming that the observed phenotypes are not an off-target effect of the shRNAs used for knockdown (Figure 6b-6e). Interestingly, the DHX9^K417R^ mutant, which lacks helicase activity due to its inability to bind ATP^52^, was also capable of suppressing PKR activation. The same was true for a construct which contained the dsRBDs of DHX9 fused to EGFP (dsRBD-EGFP). Rescue experiments in MCF-7 revealed that expression of the DHX9 dsRBD-EGFP fusion protein was sufficient to suppress not only activation of PKR, but also the RNA expression pathways described in Figure 5 and activation of RNase L (Figure 6f, g, h and I, Figure S8) These findings indicate that the DHX9 dsRBDs are likely sufficient to suppress activation of PKR, the type I IFN pathway and RNase L in the absence of ADAR1 and DHX9. However, only wild-type DHX9 could rescue the reduced foci formation observed after DHX9 knockdown (Figure 6c, e, j and k). This finding indicates that the PKR activation phenotype and the reduced proliferation phenotype are uncoupled.

**Figure 6:**
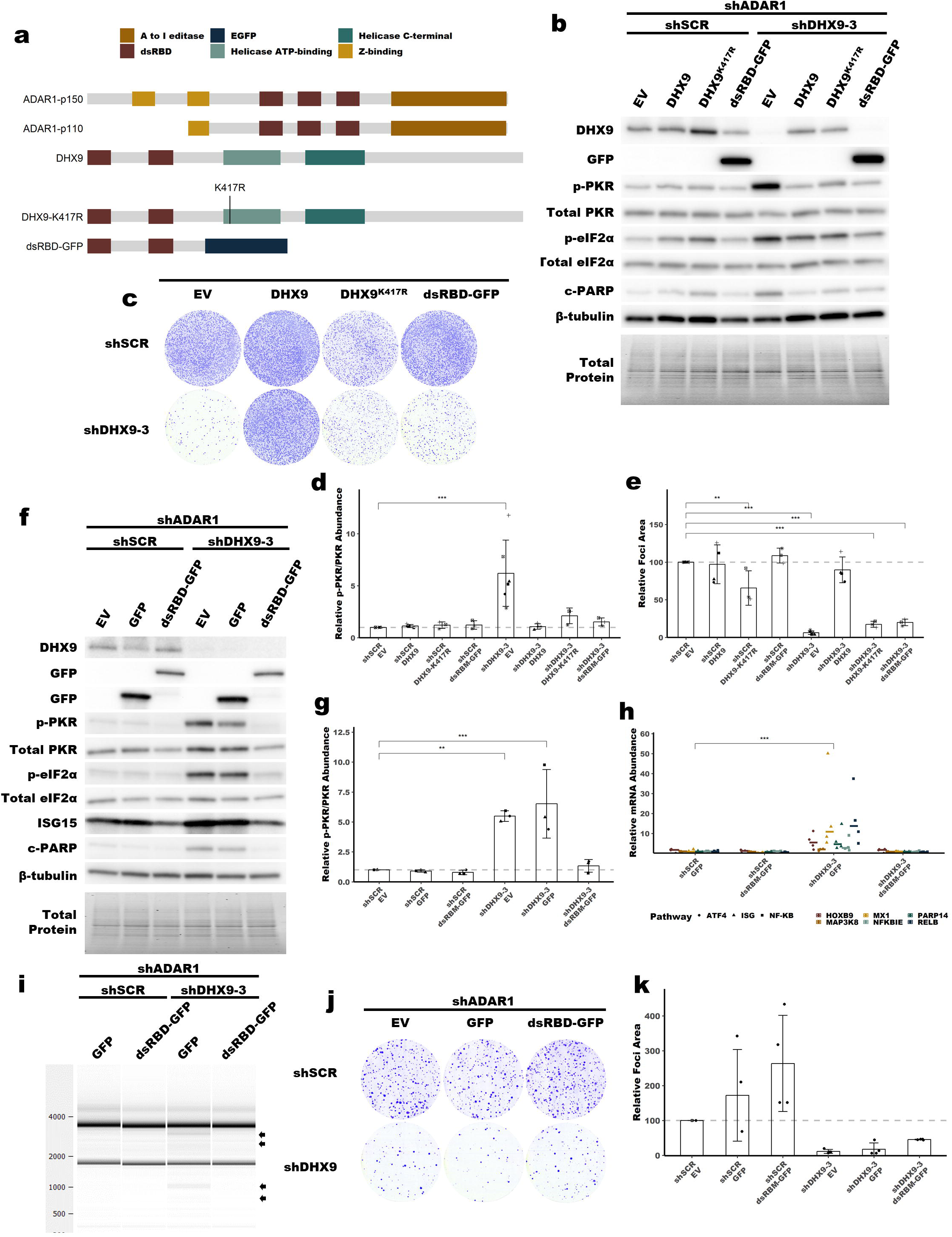
Rescue of PKR activation by ADAR1-p110, ADAR-p150, DHX9 and DHX9 mutants. **a** Schematic showing the domain structure of ADAR1 isoforms, DHX9 and mutants of DHX9, dsRBD refers to the dsRNA Binding Domain. Representative immunoblot showing the phenotype of ADAR1 and DHX9 knockdown with DHX9, DHX9^K417R^ or dsRBD-EGFP overexpression in SK-BR-3 **b**, or MCF-7 **f**. Immunoblots for other replicates and uncropped blots can be found in Source Data Figures. Representative foci formation phenotype of ADAR1 and DHX9 knockdown with DHX9, DHX9^K417R^ or dsRBD-EGFP overexpression in SK-BR-3 **c**, or MCF-7 **j**. Fold change of PKR phosphorylation at Thr-446 upon ADAR1 and DHX9 knockdown with DHX9, DHX9^K417R^ or dsRBD-EGFP overexpression in SK-BR-3 **d**, or MCF-7 **g**, quantified from immunoblots in **b** and **f** respectively. Quantification of protein expression for other proteins of interest can be found in Figure S7b-f and S8b-g. Quantification of relative foci area is shown in **e** SK-BR-3 and **k** for MCF-7. **h** RNA expression of representative genes from upregulated pathways in Figure 5 as determined by qRT-PCR for the MCF-7 rescue experiment. Statistical analysis for each pathway is shown in Figure S8h-j. **i** Analysis of rRNA integrity upon knockdown of ADAR1 and DHX9 in MCF-7 with overexpression of EGFP or dsRBD-EGFP. Arrows indicate canonical RNase L cleavage products^50^. Bars represent the average of at least three replicates, error bars are +/− SD. * p <0.05, ** p <0.01, *** p < 0.001. P-values determined by Dunnett’s test.

Consistent with the observation that the DHX9 dsRBDs are sufficient to suppress PKR activation, knockdown of DDX17, which lacks dsRBDs, did not cause substantial PKR activation in SK-BR-3 (Figure S9). Unlike DHX9, knockdown of DDX17 had no effect on cell proliferation as measured by the foci formation assay (Figure S9b and S9c). While combined knockdown of DHX9 and ADAR1 in SK-BR-3 caused a 5-10 fold increase in PKR phosphorylation, the combined knockdown of DDX17 and ADAR1 caused only a modest 1.5-fold increase in PKR phosphorylation (Figure S9a and S9d). This finding underscores the novel role of the DHX9 helicase and its dsRBD, a unique domain among this large family of RNA helicases.

### The p110 and p150 isoforms of ADAR1 suppress PKR activation in the absence of DHX9

Next, we turned to ADAR1, and asked which isoform of ADAR1 is sufficient to suppress PKR activation in the absence of DHX9. We used the same approach as above, a rescue experiment with overexpression of ADAR1-p110 or ADAR1-p150. Interestingly, we found that both ADAR1 isoforms were sufficient to suppress PKR activation upon loss of DHX9 (Figure 7a and c, Figure S10). However, overexpression of neither ADAR1 isoform was able to rescue the foci formation phenotype, again indicating that the PKR activation and cell proliferation phenotypes are uncoupled (Figure 7b and 7d).

**Figure 7:**
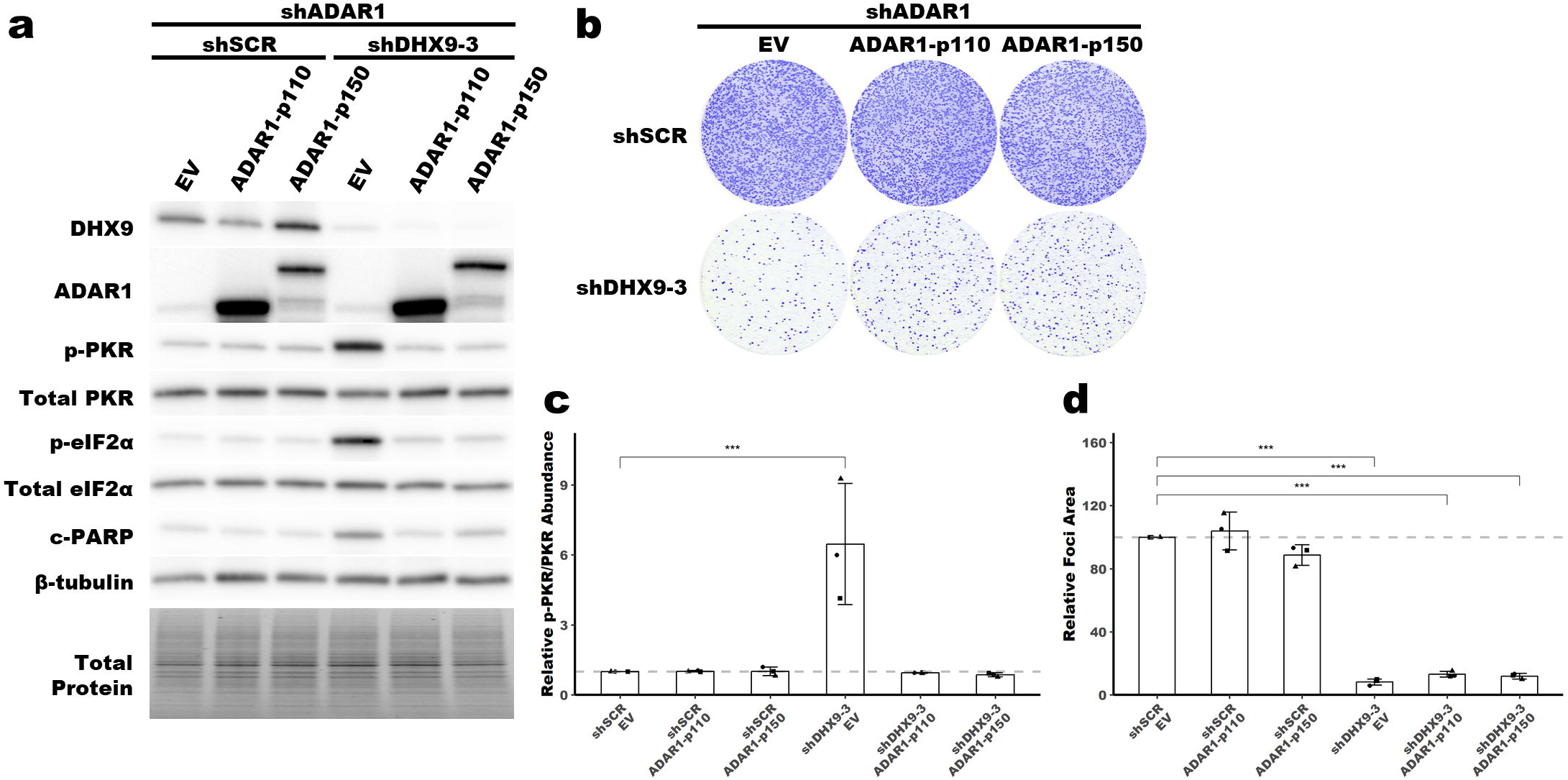
Rescue of PKR activation by ADAR1-p110 and ADAR-p150 Representative immunoblot showing the phenotype of ADAR1 and DHX9 knockdown with ADAR1-p110 or ADAR1-p150 overexpression in SK-BR-3. **a**. Immunoblots for other replicates and uncropped blots can be found in Source Data Figures. Representative foci formation phenotype of ADAR1 and DHX9 knockdown ADAR1 isoform overexpression **b**. Quantification of relative foci area is shown in **d**. Fold change of PKR phosphorylation at Thr-446 upon ADAR1 and DHX9 knockdown with ADAR1 isoform overexpression **c**, quantified from immunoblots in **a**. Quantification of protein expression for other proteins of interest can be found in Figure S10a-f. Bars represent the average of at least three replicates, error bars are +/− SD. *** p < 0.001. P-values determined by Dunnett’s test.

## Discussion

In recent years, ADAR1 has become an important therapeutic target for breast and other cancers. It is clear from the literature that depletion of ADAR1 in ADAR1-dependent cell lines leads to activation of dsRNA sensors and innate immunity programs that lead to cell death^22,34,35^. Yet unclear is what distinguishes ADAR1-dependent cell lines from ADAR1-independent cell lines – those that are insensitive to ADAR1 depletion. Elevated ISG expression has been proposed as a potential prerequisite for ADAR1-dependency, but some ADAR1-independent cell lines exhibit elevated ISG expression^22,35^. As such, more information is needed to identify the factors that establish ADAR1-dependency or ADAR1-independency. To begin to fill in some of the knowledge gaps surrounding ADAR1, we utilized proximity labeling to identify putative ADAR1 interacting proteins, specifically focusing on the less studied ADAR1-p110 isoform.

Of the proteins identified by proximity labeling, the DEAH box helicase DHX9 was of particular interest. Like ADAR1 and PKR, DHX9 possesses dsRBDs, a singularly unique trait among the DEAD/DEAH box RNA helicase family. Like many RNA helicases, DHX9 is important in numerous cellular processes, ranging from mRNA processing to resolution of R-Loops^53,54^, and has been shown to promote antiviral immunity by activation of the type I IFN, though those studies were not performed using cancer cell lines^55,56^. DHX9 expression is strongly correlated with ADAR1-p110 expression in breast cancer. Both genes are located on the q-arm of chromosome 1, though they are separated by 28 mb and are thus unlikely to be physically co-regulated. Consistent with other reports in the literature, we show here that ADAR1 and DHX9 likely interact directly^40^. While future experiments are needed to verify that ADAR1 interacts directly with DHX9, the data described here support an RNA-independent interaction between ADAR1-p110 and DHX9, and an RNA-dependent interaction between ADAR1-p150 and DHX9. This result contradicts previous findings, in which ADAR1-p150 and DHX9 co-immunoprecipitated after RNase A treatment^48^. This discrepancy may be due to different cell lines used. The importance of the ADAR1-DHX9 interaction is unclear from this work. Given that knockdown of either DHX9 or ADAR1 alone can induce PKR activation in ADAR-dependent cell lines, and that knockdown of both proteins is required to do the same in ADAR-independent cell lines, it seems unlikely that the interaction between ADAR1 and DHX9 is required for suppression of dsRNA sensing. That of course does not preclude other functions for a DHX9-ADAR1 complex, possibly in resolution of R-loops as both proteins have been implicated in R-loop homeostasis^29,53^. More studies are needed to structurally assess the interaction and directly perturb the interaction to understand what function it may have.

Here we report that in addition to being an essential gene in breast cancer, DHX9 suppresses dsRNA sensing. In ADAR1-dependent cell lines, knockdown of DHX9 alone – much like knockdown of ADAR1 alone – caused activation of the dsRNA sensor PKR^35^. Like ADAR1 knockdown, DHX9 knockdown had no effect on PKR activation in ADAR1-independent cell lines. However, combined knockdown of DHX9 and ADAR1 caused robust activation of PKR in those cells. This finding indicates that ADAR1 and DHX9 function redundantly to suppress PKR activation in ADAR1-independent cell lines and provides an explanation for why PKR is not activated in ADAR1-independent cells upon ADAR1 knockdown. In addition to suppression of PKR activation, we also observed that DHX9 and ADAR1 redundantly suppress activation of several other dsRNA sensing pathways in MCF-7. Knockdown of both proteins caused activation of IFN-I signaling, likely via MDA5 activation, as previously shown for ADAR1^13,15^. We also observed activation of OAS-RNaseL, and increased expression of the ATF4 and NF-KB targets, likely downstream of PKR activation^57–59^. Taken together, combined knockdown of DHX9 and ADAR1 in MCF-7 creates a viral mimicry phenotype, where multiple innate immunity pathways against RNA viruses have been activated. In contrast to MCF-7, we only observed activation of PKR in other cell lines after either DHX9 knockdown alone or combined knockdown with ADAR1. Transfection of the same cell lines with poly(I:C) revealed that each cell line was capable of activating PKR, the type I IFN pathway and RNase L (Figures S6h and 4k), which suggests that other variables are preventing activation of those pathways upon loss of DHX9 and ADAR1 in unresponsive cell lines. Given that PKR can be activated by much shorter dsRNAs (>30 bp^60^) than MDA5 (>500 bp^61^), it is possible that a different set of RNAs is responsible for activation of each pathway. Further studies are needed to identify other proteins that function to suppress dsRNA sensing, and to identify the dsRNAs that are activating the various pathways.

Rescue experiments revealed that the helicase activity of DHX9 was dispensable for suppression of PKR activation, and that the N-terminal dsRBDs of DHX9 were sufficient to suppress PKR activation in the absence of ADAR1 in ADAR1-independent cell lines. These data are consistent with a model in which DHX9 competes with PKR for dsRNA binding. This competition could come in the form of DHX9 binding directly to PKR and preventing its binding to dsRNA, or DHX9 binding dsRNA and preventing PKR binding. While DHX9 and PKR interact, and in fact DHX9 is phosphorylated by PKR ^62^, it is our opinion that a protein-protein interaction model is less likely considering that DHX9 is nuclear localized whereas PKR is largely localized to the cytoplasm ^54,63–65^. Conversely, DHX9 may compete with PKR for binding of dsRNA and do so by sequestering some dsRNAs within the nucleus. Sequestration of some dsRNAs within the nucleus is an important means of preventing dsRNA sensors activation. During mitosis, nuclear dsRNAs diffuse into the cytosol and activate PKR^66^. Interestingly, we did not observe a global shift in dsRNA localization following knockdown of DHX9 and ADAR1 in MCF-7 (Figure S11e-f). This finding may suggest that only a small subset of dsRNAs are responsible for activation of dsRNA sensors following loss of ADAR1 and DHX9 in those cells. This hypothesis is consistent with previous studies of ADAR1 depletion in which it was observed that only 2% of A-to-I edits are required to prevent MDA5 activation^67^. The hypothesis that only a subset of endogenous dsRNAs is responsible for activation PKR or MDA5 following loss of ADAR1 and/or DHX9 could also explain the varied results seen here across different cell lines. Some cell lines may be more sensitive to loss of ADAR1 and/or DHX9 based on the expression of specific endogenous dsRNAs.

Previously, the ADAR1 p150 isoform, and not the p110 isoform, was shown to be responsible for suppression of PKR activation^19,34^. Through rescue experiments, we show here that both isoforms are sufficient to suppress PKR activation in the absence of DHX9 in ADAR1-independent cell lines. It is important to note that the role of ADAR1-p110 in suppressing PKR activation has likely been missed previously due to the expression of DHX9. The finding that ADAR1-p110 can suppress PKR activation in the absence DHX9 highlights a redundant role of these nuclear dsRNA binding proteins. An article that was published during the preparation of this manuscript showed that the dsRBDs of ADAR1, ADAR2, and STAU1 were sufficient to suppress PKR activation^68^. Additionally, we and others have previously shown that the phenotype of ADAR1 depletion in ADAR1-dependent cells could at least be partially rescued by overexpression of an editing deficient ADAR1^34,35^. Based on these findings and our rescue experiment with the DHX9 dsRBDs, it is likely that ADAR1-p150 and ADAR1-p110 suppress dsRNA sensing by competing with PKR for dsRNA binding. For the ADAR1-p150 isoform, this competition with PKR for dsRNA binding is likely to be direct due to the cytoplasmic localization of ADAR1-p150 and PKR. However, for the nuclear localized ADAR1-p110 (Figure S11d), we propose that like DHX9, ADAR1-p110 may function to sequester PKR activating dsRNAs in the nucleus. Interestingly, the DHX9-dsRBD-EGFP fusion protein, which we showed was capable of suppressing PKR activation, localizes to the cytoplasm (Figure S11c). Together these findings suggest that there may be multiple mechanisms to prevent PKR activation – direct competition, like ADAR1-p150; and sequestration of dsRNAs, as may be the case for ADAR1-p110 and DHX9.

Our rescue experiments revealed that while the helicase activity of DHX9 was dispensable for suppression of dsRNA sensing, it was required for cell viability. Given that DHX9 has been shown to have an important role in many cellular processes, ranging from processing of mRNAs to resolution of R-Loops, we suspect the reduced viability associated with loss of DHX9 helicase activity is related to one or more of these additional DHX9 roles^53,69^

Induction of viral mimicry has great potential as a therapeutic approach for multiple cancers, including TNBC^70–73^. In addition to the cell intrinsic effects of activating innate immune pathways within the tumor, the signaling that occurs after activation of those pathways can promote anti-tumor immunity, especially when combined with checkpoint inhibitors^74–76^. Combined therapies targeting ADAR1 and DHX9 may serve as an effective means of treating breast and potentially other cancers by inducing viral mimicry.

## Materials and Methods

### Cell culture

Breast cancer cell lines (MCF-7 (RRID:CVCL_0031), SK-BR-3 (RRID:CVCL_0033), BT-549 (RRID: CVCL_1092), MDA-MB-231 (RRID: CVCL_0062), HCC1806 (RRID: CVCL_1258), MDA-MB-468 (RRID: CVCL_0063) and 293T (RRID: CVCL_0063) were obtained from American Type Culture Collection. All cell lines were cultured in Dulbecco’s modified Eagle’s medium (DMEM) (Hyclone) with 10% fetal bovine serum (BioTechne), 2 mM glutamine (Hyclone), 0.1 mM nonessential amino acids (Hyclone), and 1 mM sodium pyruvate (Hyclone).

### Viral Production and Transduction

Lentivirus was produced by Turbo DNAfection 3000 or LipoFexin (Lamda Biotech) transfection of 293T cells with pCMV-VSV-G, pCMV-ΔR8.2, and the appropriate plasmid for expression of genes of interest or shRNAs. Virus was harvested 48 hours post-transfection. Cells were transduced with lentivirus for 16 hours in the presence of 10 µg/mL protamine sulfate (Sigma-Aldrich). The cells were selected with puromycin at 2 µg/mL (Sigma-Aldrich), 150 μg/mL hygromycin (Invitrogen) or 10 µg/mL Blasticidin (Invitrogen).

### Plasmids

APEX2 was PCR amplified from pcDNA3-APEX2-NES, a kind gift from the laboratory of Kendall Blumer at Washington University in St. Louis. ADAR1-p110 was PCR amplified from pLVX-p110-ADAR1 described previously^35^. APEX2 and p110 were cloned into pLVX-IRES-puro (Takara, 632183) via a series of restriction enzyme digests and ligations. The final plasmids pLVX-3xFLAG-APEX2 and pLVX-3xFLAG-APEX2-linker-p110 were confirmed by digestion and sequencing. The linker consists of three repeats of Gly-Gly-Gly-Gly-Ser. Lentiviral shRNA constructs in the pLKO.1-puro vector were purchased as glycerol stocks from Millipore Sigma. For shADAR1, the shRNA was subcloned into pLKO.1-hygro, a gift from Bob Weinberg (Addgene, #24150). The sequences for the shRNA-scramble (shSCR) and shRNA-ADAR1 (shADAR1) were described and validated previously^35^. The sequences for shRNAs against DHX9 and DDX17 are in Supplementary Table 1. For sgDHX9, oligos encoding the sgRNAs (Supplemental Table 1) were cloned into lentiCRISPR v2, a gift from Feng Zhang (Addgene, #52961)^77^. Overexpression constructs for DHX9 and ADAR were generated by PCR amplification and ligation into pLV-EF1a-IRES-Blast vector, a gift from Tobias Meyer (Addgene, #85133)^78^. For DHX9 overexpression, wobble mutants were made to reduce shRNA targeting. Mutagenesis primers for DHX9 K417R and shRNA-resistant codons (designed using the Synonymous Mutation Generator^79^) are included in Supplemental Table 1. The DHX9-dsRBD-EGFP construct was generated by digestion of pLV-EF1-DHX9 with SpeI and EcoRI and ligation of EGFP in place of the 3’ portion of DHX9. The resulting construct codes for the first 344 amino acids of DHX9 fused to EGFP.

### Immunoblot

Cell pellets were lysed and sonicated in RIPA Buffer (50 mM Tris pH 7.4 (Ambion), 150 mM NaCl (Ambion), 1% Triton X-100 (Sigma-Aldrich), 0.1% sodium dodecyl sulfate (Promega) and 0.5% sodium deoxycholate (Sigma-Aldrich) with 1x HALT Protease Inhibitor (Pierce). Protein was quantified using the DC Assay kit (Bio-Rad) and diluted in SDS Sample Buffer (125 mM Tris pH 6.8, 30% glycerol, 10% sodium dodecyl sulfate, 0.012% bromophenol blue) prior to denaturation by heating to 95 °C for 7 minutes. Thirty micrograms of protein lysate were resolved on 4-12% TGX Acrylamide Stain-Free gels (Bio-Rad). Stain-Free gels were imaged prior to transfer to PVDF membrane (Millipore or Bio-Rad) by TransBlot Turbo (Bio-Rad). The blots were then probed with the appropriate primary antibodies: ADAR1 (Santa Cruz, sc-73408; Bethyl Laboratories, A303-883A), DDX17 (Thermo Scientific, PA5-84585), DHX9 (Bethyl, A300-855A), DDX54 (Novus Biologicals, NB100-60678), eIF2a (Abcam, ab5369), eIF2a-Ser-51-P (Abcam, ab32157), Fibrillarin (Santa Cruz, sc-25397), beta-tubulin (Abcam, ab6046), ISG15 (Santa Cruz, sc-166755), cleaved PARP (Cell Signaling, #9541), PKR (Cell Signaling, #3072), PKR Thr-446-P (Abcam, ab32036), MDA5 (Cell Signaling, #5321S), MAVS (Cell Signaling, #3993S). Primary antibodies were detected with horseradish-peroxidase conjugated secondary antibodies (Jackson ImmunoResearch) and detection was carried out with Clarity Western ECL Substrate (Bio-Rad). Densitometry was performed using Image Lab (Bio-Rad). Band intensity was normalized to total protein measured by imaging of the Stain-Free gel.

### Proximity Labeling by APEX2

SK-BR-3, MCF-7, or MDA-MB-231 cells expressing pLVX-puro-FLAG-APEX2 or pLVX-puro-FLAG-APEX2-ADAR1p110 were grown to ∼80% confluency in a 15 cm dish. Quencher solution (10 mM sodium azide, 10 mM sodium ascorbate, and 5 mM Trolox (Sigma-Aldrich) in 1X PBS) was prepared at 1X and 2X concentrations. Prior to labeling, cells were incubated in 10 mL of culture media containing 500 μM biotin phenol (Toronto Research Chemicals, B397770) for 30 min at 37 °C. Next, hydrogen peroxide was added to the cells at 1 mM and incubated at room temperature for 1 min. Immediately one volume of 2X quencher solution was added to the cells to stop labeling. The cells were washed twice with 1X quencher solution. Cells were harvested by scraping in 1X quencher solution and lysed in RIPA buffer containing 10 mM sodium azide, 10 mM sodium ascorbate, 5 mM Trolox and 1x HALT. Biotin labeling was verified by immunoblotting with HRP-streptavidin (Abcam, ab7403). Biotinylated proteins were purified using streptavidin magnetic beads (Thermo Fisher Scientific, 88816). Streptavidin magnetic beads were washed twice with RIPA containing HALT and quenching agents. The lysate from above was incubated with the beads for 1 hour at room temperature. The beads were washed in the following order: once with RIPA containing HALT and quenching agents, once with RIPA, once with 1 M KCl, once with 2 M urea pH 8.0, twice in RIPA and once with water. Elution was performed in 1X SDS-Sample buffer by heating at 95 °C for 10 minutes. The eluate was analyzed by LC-MS-MS, see below.

### Mass Spectrometry

Liquid-chromatography and tandem mass spectrometry was performed by MSBioWorks (Ann Arbor, MI). The eluates from the streptavidin pulldown above were processed by SDS-PAGE using 10% Bis-Tris NuPage Mini-gel (Invitrogen) with the MES buffer system. The gel was run 2cm. The mobility region was excised and processed by in-gel digestion with trypsin using a robot (ProGest, DigiLab). For the trypsin digestion, the gel slices were washed with 25mM ammonium bicarbonate followed by acetonitrile. The samples were reduced with 10mM dithiothreitol at 60°C followed by alkylation with 50mM iodoacetamide at room temperature. Subsequently proteins were digested with trypsin (Promega) at 37°C for 4 h. The trypsin digestion was quenched with formic acid and the supernatant was analyzed directly without further processing. The digested sample was analyzed by nano LC-MS/MS with a Waters M-Class HPLC system interfaced to a ThermoFisher Fusion Lumos mass spectrometer. Peptides were loaded on a trapping column and eluted over a 75 µm analytical column at 350 nL/min; both columns were packed with Luna C18 resin (Phenomenex). The mass spectrometer was operated in data-dependent mode, with the Orbitrap operating at 60,000 FWHM and 15,000 FWHM for MS and MS/MS respectively. APD was enabled and the instrument was run with a 3 s cycle for MS and MS/MS. Five hours of instrument time was used for the analysis of each sample.

### Analysis of Mass Spectrometry Data

Data were searched using a local copy of Mascot (Matrix Science) with the following parameters: Enzyme – Trypsin/P; Database – SwissProt Human (concatenated forward and reverse plus common contaminants); Fixed modification – Carbamidomethyl; Variable modifications – Oxidation; Acetyl; Pyro-Glu; Deamidation; Mass values – Monoisotopic; Peptide Mass Tolerance – 10 ppm; Fragment Mass Tolerance – 0.02 Da; Max Missed Cleavages – 2. Mascot DAT files were parsed into Scaffold (Proteome Software) for validation, filtering and to create a non-redundant list per sample. Data were filtered using a 1% protein and peptide FDR and requiring at least two unique peptides per protein. Fold change of protein abundance was determined by DESeq2 using spectral counts (see Data Availability below for scripts). Overrepresentation analysis was performed using ‘enrichR’ in R^80^. The cutoff used for enrichment for the overrepresentation analysis was an FDR < 0.05 and a log2 fold change of > 0.5.

### Immunoprecipitation

Cell lysates prepared in RIPA with 1X HALT. RNaseOUT (Thermo Fisher) RNase inhibitor was added to the lysis buffer at 0.5 U/μL when RNase A was not used. One milligram of protein lysate was mixed with 2-10 µg of IgG or specific antibody overnight at 4 °C with rotation. For samples treated with RNase A, 20 μg of RNase A (Invitrogen) was added to the lysate during overnight mixing with the antibody. Protein G Dynabeads (Thermo Fisher, 25 μL per sample) were prepared by washing twice in the lysis buffer. Prepared beads were mixed with lysates for 30 min at 4 °C with rotation. Supernatants were collected and beads were washed three times in the lysis buffer, and eluted by mixing the beads in SDS sample buffer and incubating at 95 °C for 7 min. Antibodies: Rabbit IgG (Jackson ImmunoResearch, 011-000-003), Mouse IgG (Jackson ImmunoResearch, 015-000-003), DHX9 (Bethyl, A300-855A), PARP (Cell Signaling, 9532S), XRN2 (Novus, NB100-57541), DDX54 (Novus, NB100-60678), DDX17 (Thermo Scientific, PA5-84585).

### Immunofluorescence

Cells were plated on glass coverslips (Corning) two days prior to fixation for immunofluorescence. Cells were washed in PBS prior to fixation with 4% paraformaldehyde (Thermo Scientific) and permeabilization with 0.15% Triton-X100 in PBS. Following permeabilization the cells were washed three times with PBS then blocked with Protein Block (Aligent/Dako, X090930-2). Primary antibodies (ADAR1 (Santa Cruz, sc-73408), Fibrillarin (Santa Cruz, sc-25397), DDX54 (Novus Biologicals, NB100-60678) or DDX17 (Thermo Scientific, PA5-84585), DHX9 (Bethyl, A300-855A), ADAR1-p150 (Abcam, Ab126745), dsRNA-J2 (Millipore-Sigma, MABE1134) and secondary antibodies (Thermo Scientific, A21207, A21203, A21202, A21206) were diluted in Antibody Diluent (Agilent/Dako, S302283-2). Antibody binding was performed in a humidity chamber for 1.5 hours for primaries and 30 minutes for secondaries. Between primary and secondary antibodies, and after secondary antibody binding the coverslips were washed in PBS. The coverslips were washed once in water before mounting on glass slides with Vectashield Antifade Mounting Media with DAPI (Vector Laboratories, H-1200-10). Fluorescence microscopy images were obtained with an Eclipse 90i microscope (Nikon) using a Plan Apochromatic 20x/NA 0.75 objective (Nikon) and a CoolSNAP ES2 monochrome digital camera cooled to 0°C (Photometrics). Fluorescence images were captured with MetaMorph version 7.8.0.0 software (Molecular Devices) and resized and formatted with Fiji.

### Transfection of poly(I:C)

The cell line indicated was transfected with high-molecular weight poly(I:C) (Invivogen) with Lipofectamine LTX (Invitrogen). Three microliters of Lipofectamine LTX was used per microgram of poly(I:C). Sixteen to 24 hours after transfection, cells were harvested in RIPA with 1X HALT or the RNA lysis buffer from the Nucleospin RNA kit (Macherey-Nagel).

### RNA Purification and RNA sequencing

RNA-sequencing was performed for two replicates of ADAR1 and/or DHX9 knockdown in MCF-7 and SK-BR-3. RNA was purified using the Nucleospin RNA kit (Macherey-Nagel). Assessment of rRNA integrity and RNA-sequencing was performed by the Genome Technology Access Center at Washington University in St. Louis. Total RNA integrity was determined using Agilent TapeStation 4200. Library preparation was performed with 500 ng to 1 ug of total RNA. Ribosomal RNA was removed by an RNase-H method using RiboErase kit (Kapa Biosystems). After rRNA depletion, the remaining RNA was then fragmented in reverse transcriptase buffer (Life Technologies) by heating to 94 degrees for 8 minutes. The RNA was reverse transcribed to yield cDNA using SuperScript III RT and random hexamers (Life Technologies) per manufacturer’s instructions. A second strand reaction was performed with DNA Polymerase I and RNase H (Qiagen) to yield ds-cDNA. The cDNA was then blunted with T4 DNA Polymerase, Polynucleotide Kinase and Klenow DNA Polymerase (Qiagen). An A base was added to the 3’ ends with Klenow (3’-4’ exo-) (Qiagen). The processed ds-cDNA was then ligated to Illumina sequencing adapters with T4 DNA Ligase (Qiagen). Ligated fragments were then amplified for 12-15 cycles using primers incorporating unique dual index tags with VeraSeq polymerase (Qiagen). Fragments were sequenced on an Illumina NovaSeq-6000 using paired end reads extending 150 bases.

### RNA Sequencing Analysis

The Illumina bcl2fastq software was used for base calling and demultiplexing, allowing for one mismatch in the indexing read. STAR version 2.7.9a1 was used for read alignment to RNA-seq to the Ensembl GRCh38.101 primary assembly. Gene counts were determined using Subread:featureCount version 2.0.32, only uniquely aligned unambiguous reads were counted. Differential gene expression was determined using DESeq2 (see Data Availability below for scripts)^81^. The experimental design for DESeq2 analysis included an interaction term between the shRNAs used for knockdown (‘shrna1 + shrna2 + shrna1:shrna2’; where ‘shrna1’ was either shSCR or shADAR and ‘shrna2’ was either shSCR or shDHX9-3). Contrasts were used to assess differential expression after singular knockdown of either ADAR1 or DHX9. Fold changes were shrunken using the ‘apeglm’ approach from DESeq2^82^. Gene set enrichment analysis was performed using ‘clusterProfiler’^83^. For analysis of transposable element expression, ‘TEcount’ from TEtranscripts^84^ was used to determine family level counts for transposable elements using a GTF file containing transposable element information from RepeatMasker (http://www.repeatmasker.org). (see Data Availability section for more information about the GTF file).

### Foci Formation Assay

Five thousand cells were plated for each condition in a 10 cm culture dish. After 10 (BT-549, MDA-MB-MB231, HCC1806, MDA-MB-468 and SK-BR-3) to 20 (MCF-7) days the cells were washed briefly with 1x PBS prior to fixation in 100% methanol for 5 min. After drying, the cells were stained with 0.005% Crystal Violet solution containing 25% methanol (Sigma-Aldrich) prior to washing excess stain away with deionized water. The plates were scanned using an ImageScanner III (General Electric). Foci area was calculated using ImageJ.

### Quantitative PCR

Reverse transcription and qPCR were performed as described previously using iScript Supermix for cDNA synthesis and iTaq for qPCR (Bio-Rad)^85^. Fold change of RNA expression was determined using the ΔΔC_t_ method with normalization to PSMA5 and OAZ1. Primers are shown in Supplemental Table 1.

### Analysis of CCLE and TCGA data

RNA-seq normalization and calculation of z-scores was performed as described previously^35^. Molecular subtypes of breast cancer cell lines and TCGA samples were defined previously^86^. Breast cancer survival analysis was performed using the R packages RTCGA and survminer^87,88^. The DHX9 expression level used for stratification in survival analysis was determine by the surv_cutpoint function of survminer.

### Data and Code Availability

Scripts used for analysis of mass spectrometry, RNA-seq and generation of all plots are available at (https://github.com/cottrellka/Cottrell-Ryu-et-al-2023), raw sequencing data and count files were deposited at the Gene Expression Omnibus (GEO) under accession <u>GSE224677</u>. RNA-seq data for cancer cell lines (CCLE_expression_full.csv, CCLE_RNAseq_rsem_transcripts_tpm_20180929.txt) were obtained from the DepMap portal (https://depmap.org/portal/download/custom/)^89^. RNAi based dependency data for DHX9 (D2_combined_gene_dep_scores) was obtained from DepMap Portal (https://depmap.org/portal/download/custom/)^90^. RNA-seq data for TCGA BRCA samples (illuminahiseq_rnaseqv2-RSEM_genes, illuminahiseq_rnaseqv2-RSEM_isoforms_normalized) and clinical data (Merge_Clinical) were obtained from the Broad Institute FireBrowse and are available at http://firebrowse.org/. The GTF file used for TEcount (GRCh38_Ensembl_rmsk_TE.gtf) is available at https://www.dropbox.com/sh/1ppg2e0fbc64bqw/AACUXf-TA1rnBIjvykMH2Lcia?dl=0.

## Acknowledgements

The authors would like to thank Leonard B. Maggi, Jr., Raleigh Kladney, Louis Kerestes, Che-Pei Kung, and Anbrielle Blake for technical assistance, Elliot Klotz and Toni Sinnwell (Genome Technology Access Center at Washington University) for RNA sequencing, Joshua B. Rubin, Jason C. Mills, Jieya Shao, and Alessandro Vindigni for discussion and suggestions. The data described here is in part based on data generated by the TCGA Research Network: https://www.cancer.gov/tcga. This work was supported by R01CA262804 (J.D.W.), K99MD016946 and R00MD016946 (K.A.C) from the National Institutes of Health, W81XWH-18-1-0025 (J.D.W.) and W81XWH-21-10466 (J.D.W.) from the Department of Defense, and a Centene Corporation Contract (P19-00559) for the Washington University–Centene ARCH Personalized Medicine Initiative (J.D.W.). L.S.T. supported by R25HG006687 and T32GM148405.

## Author Contributions

K.A.C, S.R. and J.D.W. conceived the project. K.A.C., S.R., L.S.T., and A.M.S. performed the experiments. K.A.C., S.R., and L.S.T. performed the data analysis. K.A.C. and S.R. wrote the manuscript. All authors edited the manuscript.

