## Supplemental Figures for "Induction of viral mimicry upon loss of DHX9 and ADAR1 in breast cancer cells"

**Contents:**

Figures S1-11
Source Data Figures for main Figures and Supplemental Figures

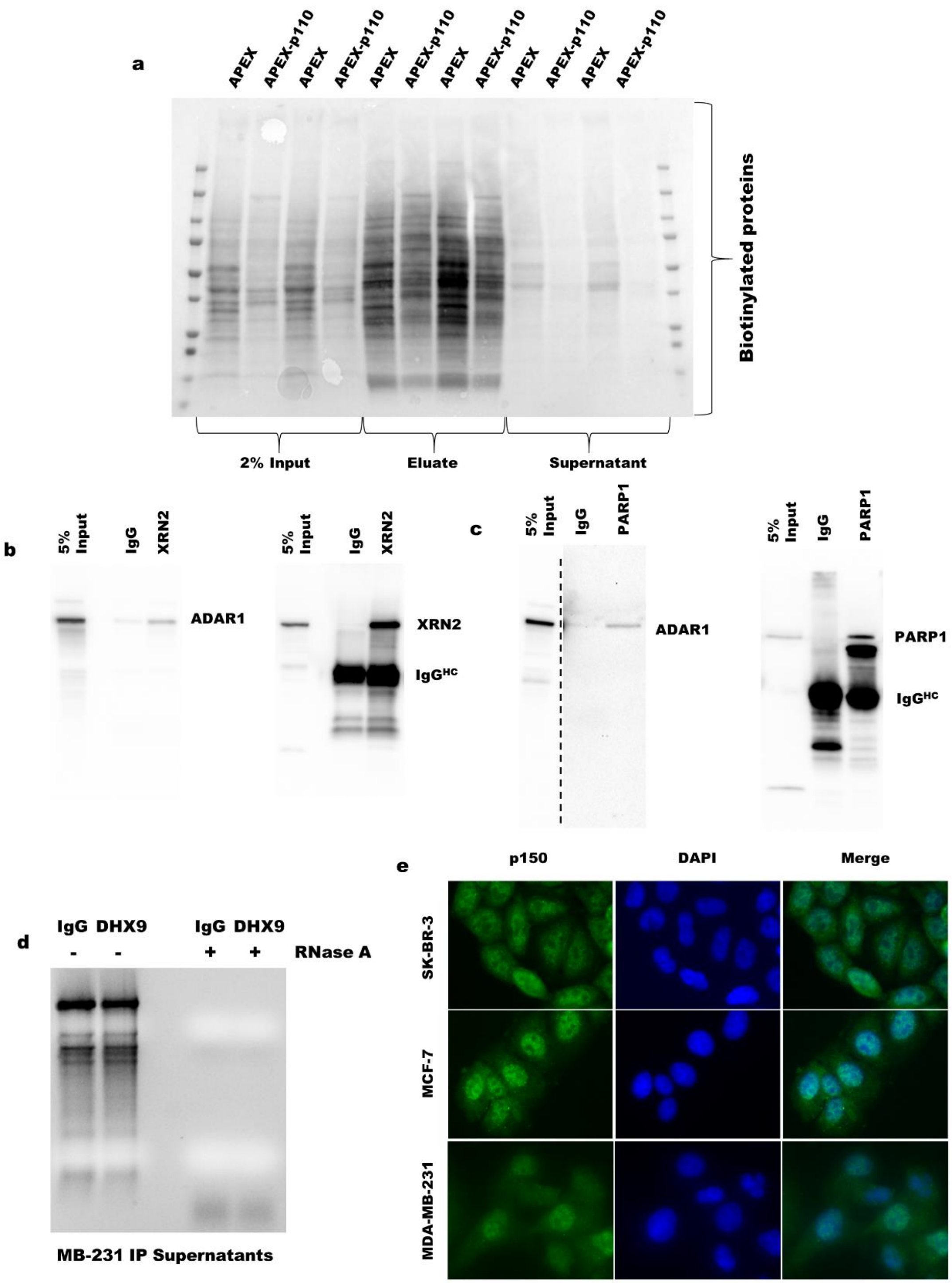


**Figure S1:**

**a** Representative streptavidin-HRP blot of the streptavidin pulldown for MCF-7. The '2% Input' represents the input used for the streptavidin pulldown following proximity labeling. The 'Eluate' is what was eluted from the streptavidin beads following pulldown. The 'Supernatant' is the supernatant following binding of the lysate to the streptavidin beads and before washing the beads. **b**-**c** Representative blots following immunoprecipitation of XRN2 or PARP1 in SK-BR-3. Input represents 5% of the lysate used for immunoprecipitation. The IgG lanes represent immunoprecipitation eluates from pulldown with anti-rabbit IgG antibody. The lanes labeled XRN2 or PARP1 indicate the eluates from immunoprecipitation with antibodies against those proteins respectively. The $IgG^{HC}$ label indicates the band corresponding to the IgG heavy chain from the antibody used for immunoprecipitation. **d** Assessment of rRNA integrity following immunoprecipitation of DHX9 in MDA-MB-231.

**Figure S1: (cont.)**

RNA was purified from the immunoprecipitation supernatants. rRNA integrity was assessed by denaturing agarose gel electrophoresis and staining with ethidium bromide. **e** Indirect immunofluorescence for ADAR1-p150 in three breast cancer cell lines.

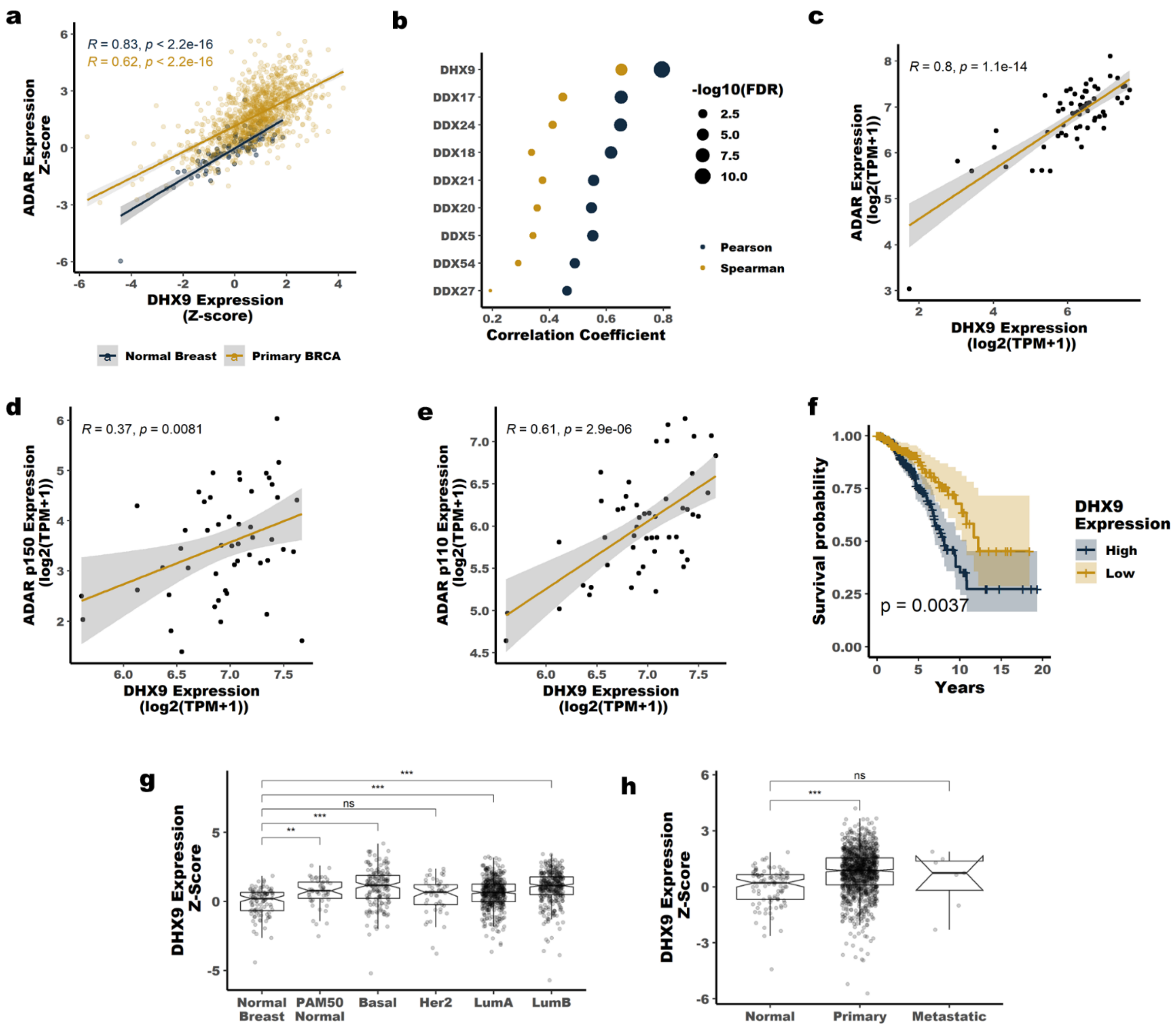


**Figure S2:**

**a** Scatter plot showing the correlation between ADAR1 and DHX9 expression at the RNA level in normal breast and primary BRCA, data from TCGA. The Pearson correlation coefficient and p-value are shown. **b** Pearson and Spearman correlation coefficients for the correlation between ADAR1 expression at the RNA level and the expression of each indicated helicase at the RNA level, data from breast cancer cell lines within CCLE. **c** Scatter plot showing the correlation between ADAR1 and DHX9 expression at the RNA level in breast cancer cell lines, data from CCLE. The Pearson correlation coefficient and p-value are shown. Scatterplots showing the correlation between ADAR1-p150 **d**, or ADAR1-p110 **e**, and DHX9 expression in breast cancer cell lines, data from CCLE. The Pearson correlation coefficient and p-value are shown. **f** Survival of breast cancer patients stratified by DHX9 expression. See methods for information on how the DHX9 expression cutoff was determined. Data from TCGA. **g** and **h** Expression of DHX9 at the RNA level based on PAM50 classification **g** or tumor site **h**, data from TCGA. * p <0.05, ** p <0.01, *** p < 0.001. P-values determined by Dunnett's test.

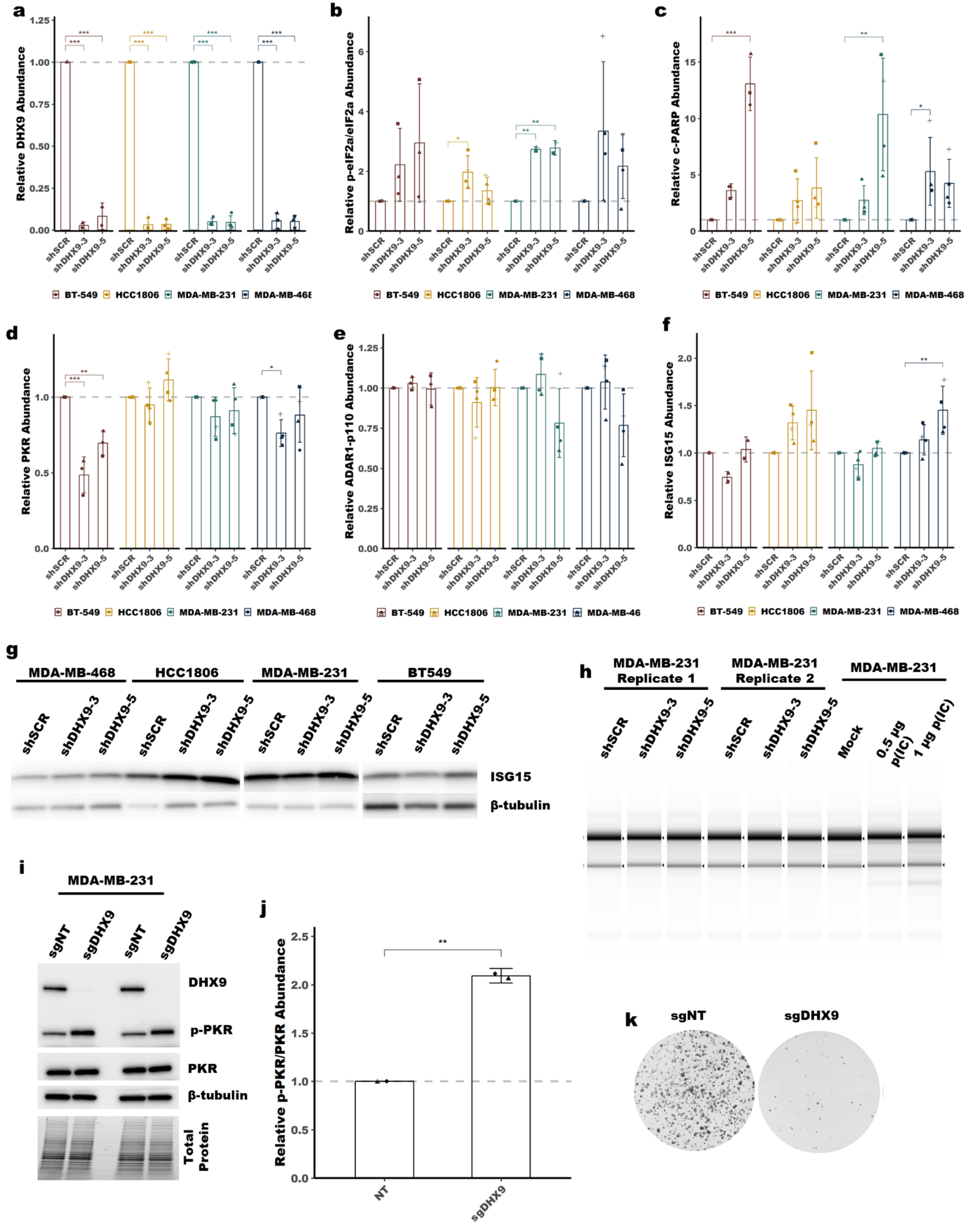
a
b
c
d
e
f
g
h
i
j
k
BT-549
HCC1806
MDA-MB-231
MDA-MB-468
shSCR
shDHX9-3
shDHX9-5
Relative DHX9 Abundance
Relative p-eIF2a/eIF2a Abundance
Relative c-PARP Abundance
Relative PKR Abundance
Relative ADAR1-p110 Abundance
Relative ISG15 Abundance
BT549
ISG15
β-tubulin
MDA-MB-231 Replicate 1
MDA-MB-231 Replicate 2
Mock
0.5 µg p(IC)
1 µg p(IC)
sgNT
sgDHX9
DHX9
p-PKR
PKR
Total Protein
Relative p-PKR/PKR Abundance
NT

Figure S3:

**Figure S3: (cont.)**

**a**-**e** Quantification of the immunoblots shown in Figure 3g. **f** Quantification of the immunoblots shown in panel **g**. **g** Representative immunoblot for ISG15 and β-tubulin in TNBC cell lines following knockdown of DHX9. Immunoblots from additional experiments can be found in the Source Data Figures. **h** Assessment of rRNA integrity in MDA-MB-231 following knockdown of DHX9 or transfection with poly(I:C) (p(I:C)). **i** Immunoblot for DHX9, p-PKR, PKR and β-tubulin following knockout of DHX9 by CRISPR-Cas9 in MDA-MB-231, two replicates are shown, quantification of the p-PKR abundance is shown in **j**. **k** Representative foci formation assay results for knockout DHX9 in MDA-MB-231. Bars represent the average of at least three replicates **a**-**e** or at least two replicates **f**, error bars are +/- SD. * $p < 0.05$, ** $p < 0.01$, *** $p < 0.001$. P-values determined by Dunnett's test.

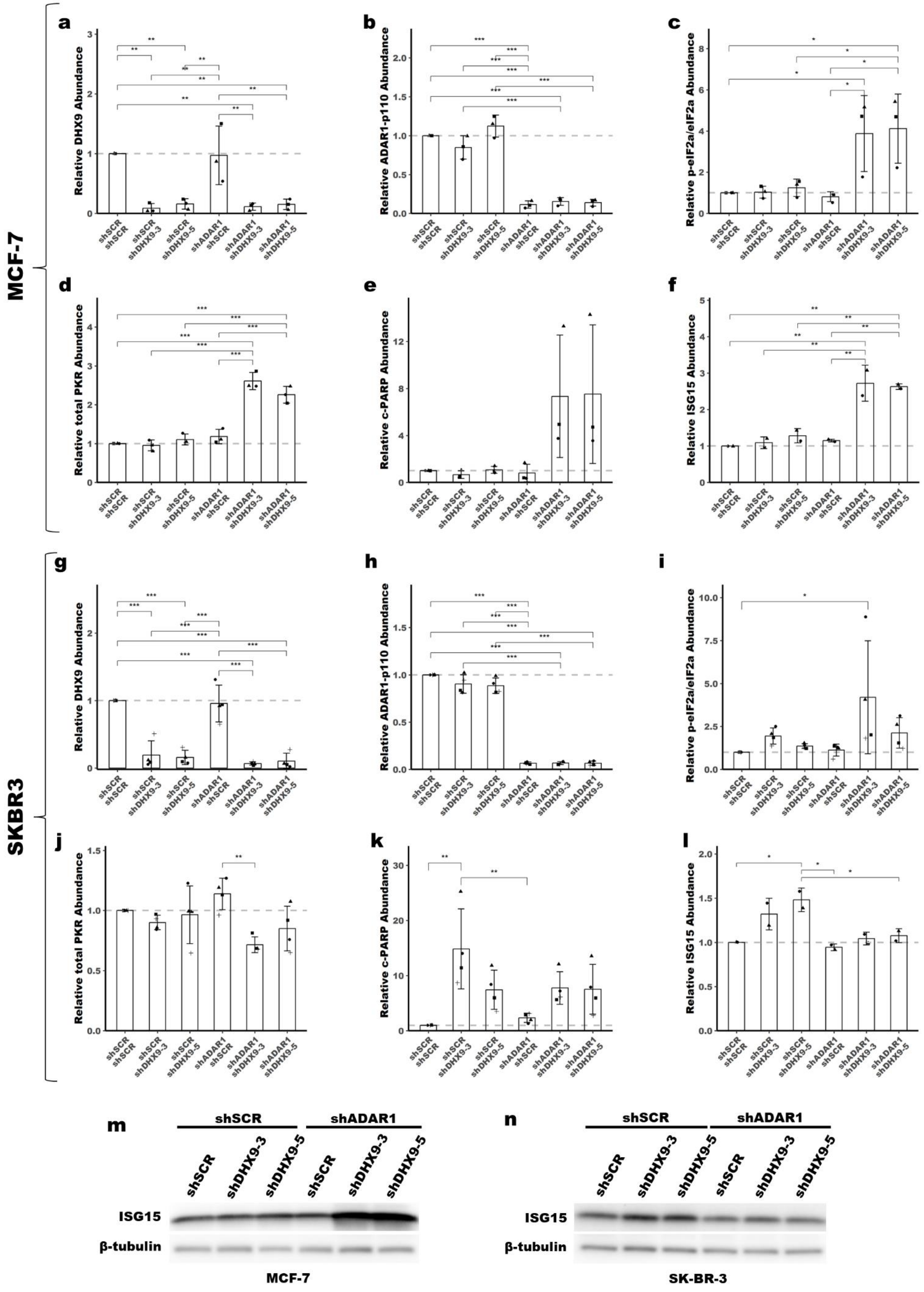
MCF-7
SKBR3
a
Relative DHX9 Abundance
b
Relative ADAR1-p110 Abundance
c
Relative p-eIF2a/eIF2a Abundance
d
Relative total PKR Abundance
e
Relative c-PARP Abundance
f
Relative ISG15 Abundance
g
Relative DHX9 Abundance
h
Relative ADAR1-p110 Abundance
i
Relative p-eIF2a/eIF2a Abundance
j
Relative total PKR Abundance
k
Relative c-PARP Abundance
l
Relative ISG15 Abundance
shSCR shSCR
shSCR shDHX9-3
shSCR shDHX9-5
shADAR1 shSCR
shADAR1 shDHX9-3
shADAR1 shDHX9-5
m
shSCR
shADAR1
shSCR
shDHX9-3
shDHX9-5
shSCR
shDHX9-3
shDHX9-5
ISG15
β-tubulin
MCF-7
n
shSCR
shADAR1
shSCR
shDHX9-3
shDHX9-5
shSCR
shDHX9-3
shDHX9-5
ISG15
β-tubulin
SK-BR-3

Figure S4:

**Figure S4: (cont.)**

**a**-**l** Quantification of the immunoblots shown in Figure 4a, 4f and panels **m** and **n** in this figure. **m**-**n** Representative immunoblot for ISG15 and β-tubulin in MCF-7 and SK-BR-3 following knockdown of DHX9 and/or ADAR1. Immunoblots from additional experiments can be found in the Source Data Figures. Bars represent the average of at least three replicates **a**-**e** and **g**-**k** or two replicates **f** and **l**, error bars are +/- SD. * $p < 0.05$, ** $p < 0.01$, *** $p < 0.001$. P-values determined by one-way ANOVA with post-hoc Tukey. Comparisons between the two different shRNAs targeting DHX9 (shDHX9-3 and shDHX9-5) were not included for clarity.

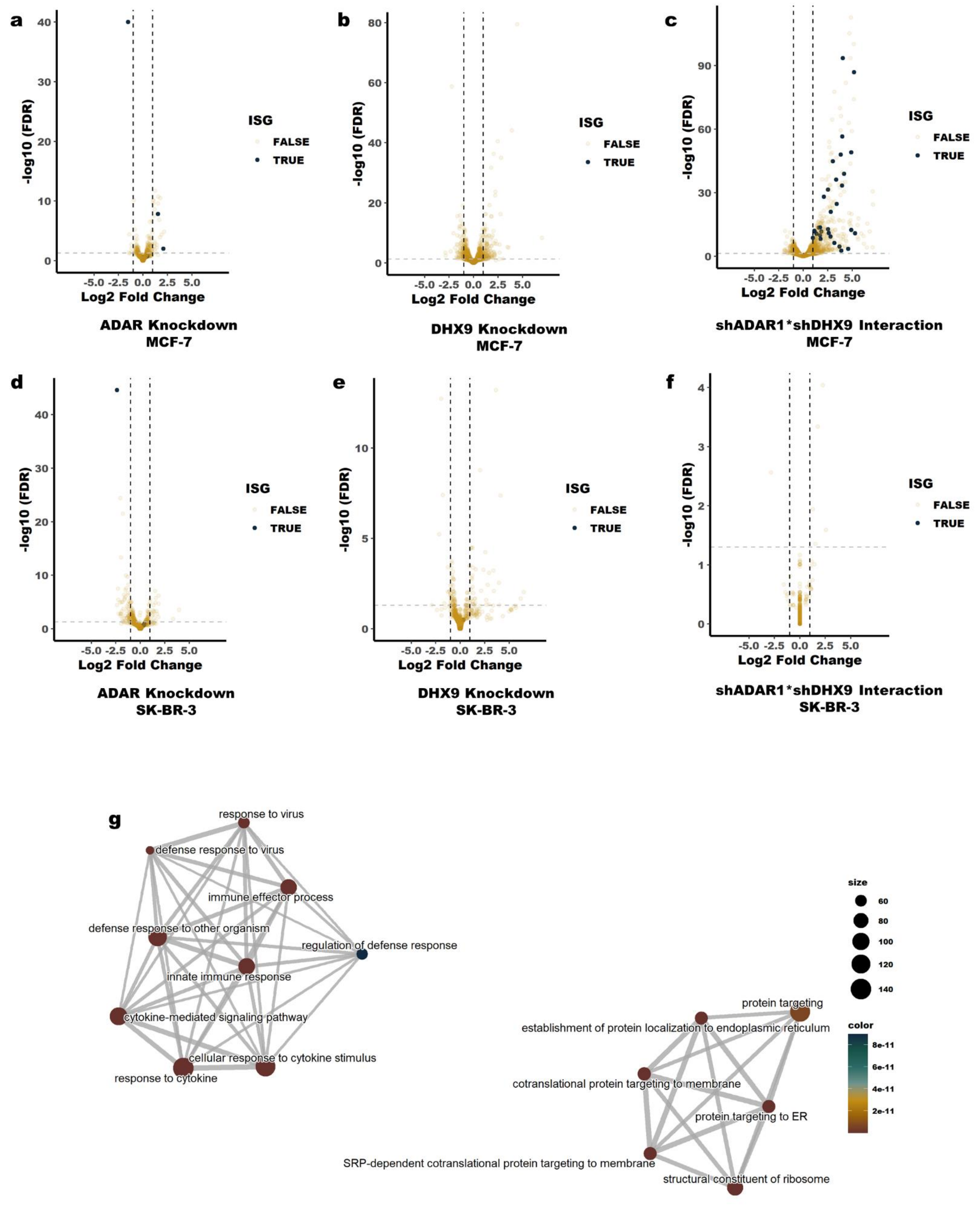
a
-log10 (FDR)
Log2 Fold Change
ISG
FALSE
TRUE
ADAR Knockdown
MCF-7
b
DHX9 Knockdown
MCF-7
c
shADAR1*shDHX9 Interaction
MCF-7
d
ADAR Knockdown
SK-BR-3
e
DHX9 Knockdown
SK-BR-3
f
shADAR1*shDHX9 Interaction
SK-BR-3
g
response to virus
defense response to virus
immune effector process
defense response to other organism
regulation of defense response
innate immune response
cytokine-mediated signaling pathway
cellular response to cytokine stimulus
response to cytokine
protein targeting
establishment of protein localization to endoplasmic reticulum
cotranslational protein targeting to membrane
protein targeting to ER
SRP-dependent cotranslational protein targeting to membrane
structural constituent of ribosome
size
60
80
100
120
140
color
8e-11
6e-11
4e-11
2e-11

**Figure S5:**

**Figure S5: (cont.)**

**a**-**f** Volcano plots showing changes in RNA expression upon knockdown of DHX9 and/or ADAR1 in MCF-7 or SK-BR-3. For panels **c** and **f**, fold-change of RNA expression was determined using an interaction term between ADAR1 and DHX9 knockdown. For all volcano plots, Core ISGs have been labeled. The ISG with lower expression upon knockdown of ADAR1 in MCF-7 and SK-BR-3 **a** and **d** is ADAR1. **g** Enrichment map for the GO terms described in Figure 5b and Supplementary Table 14 (based on combined knockdown of ADAR1 and DHX9 in MCF-7). The GO terms in the left cluster are upregulated following knockdown of DHX9 and ADAR1, while the terms in the right cluster are downregulated. The size of each point represents the number of genes associated with the GO term and the color is the FDR corrected p-value.

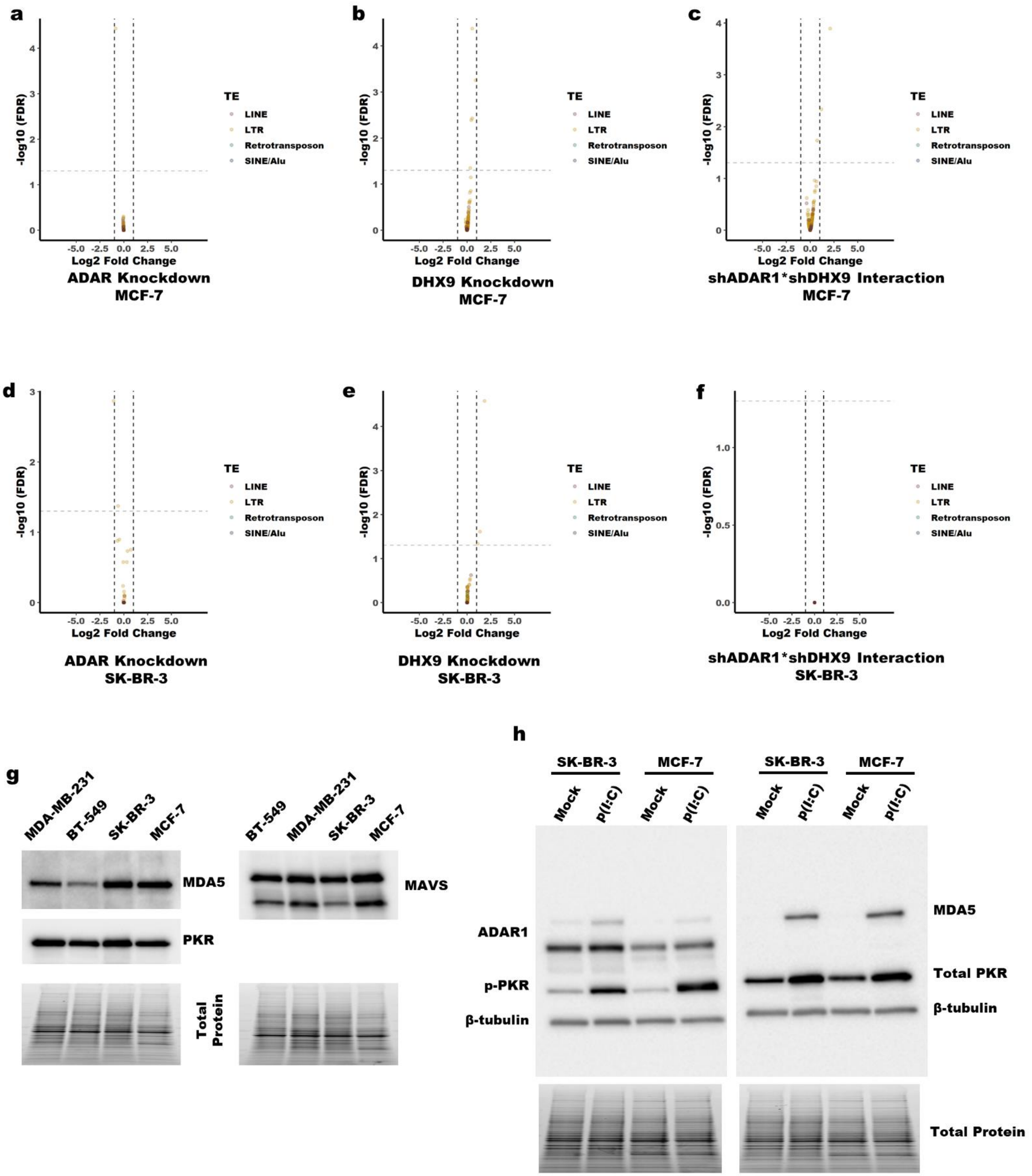


**Figure S6:**

**a**-**f** Volcano plots showing changes in transposable element expression upon knockdown of DHX9 and/or ADAR1 in MCF-7 or SK-BR-3. For panels **c** and **f**, fold-change of RNA expression was determined using an interaction term between ADAR1 and DHX9 knockdown. Individual points are colored based on the transposable element family. **g** Immunoblot for MDA5, PKR and MAVS expression in breast cancer cell lines of interest. **h** Immunoblot for assessing activation of PKR and type I IFN signaling following transfection of MCF-7 or SK-BR-3 with p(I:C).

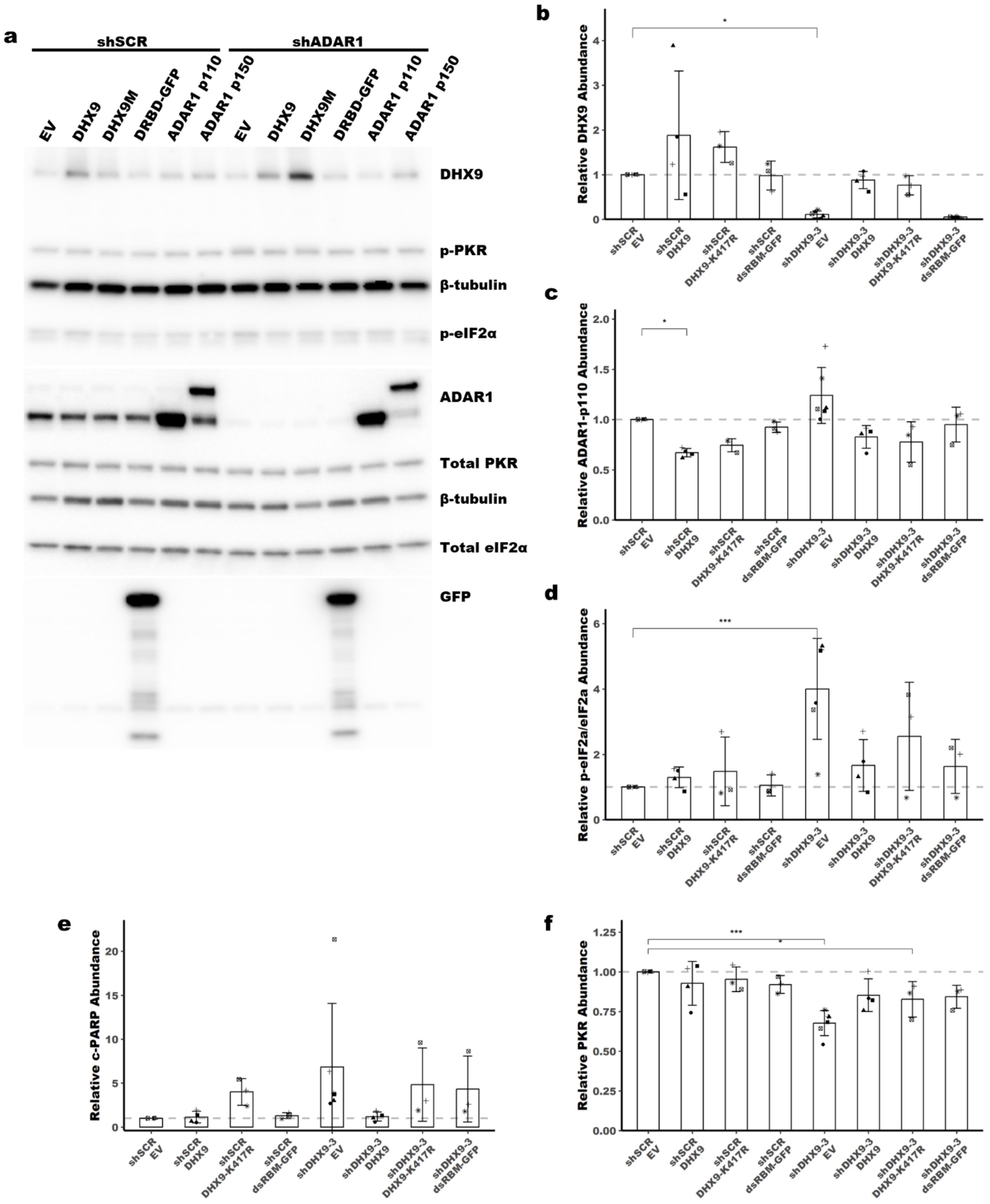


**Figure S7:**

**a** Representative immunoblot showing the expression of the constructs used in the rescue experiments described in Figure 6. **b**-**f** Quantification of the immunoblot in Figure 6b. Bars represent the average of at least three replicates, error bars are +/- SD. * $p < 0.05$, ** $p < 0.01$, *** $p < 0.001$. P-values determined by Dunnett's test.

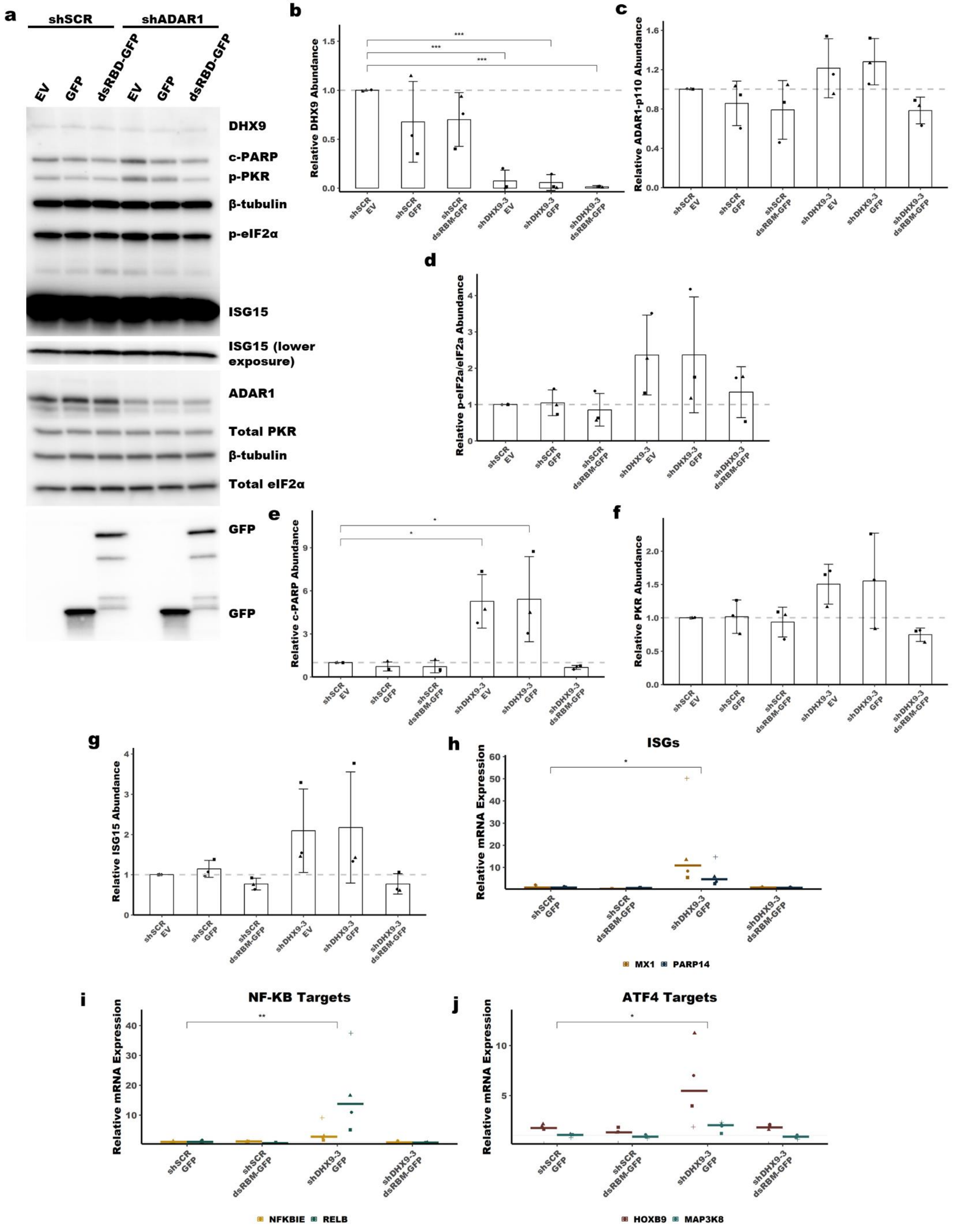
a
shSCR
shADAR1
EV
GFP
dsRBD-GFP
EV
GFP
dsRBD-GFP
DHX9
c-PARP
p-PKR
β-tubulin
p-eIF2α
ISG15
ISG15 (lower exposure)
ADAR1
Total PKR
β-tubulin
Total eIF2α
GFP
GFP
b
Relative DHX9 Abundance
c
Relative ADAR1-p110 Abundance
d
Relative p-eIF2a/eIF2a Abundance
e
Relative c-PARP Abundance
f
Relative PKR Abundance
g
Relative ISG15 Abundance
shSCR EV
shSCR GFP
shSCR dsRBM-GFP
shDHX9-3 EV
shDHX9-3 GFP
shDHX9-3 dsRBM-GFP
h
ISGs
Relative mRNA Expression
MX1
PARP14
i
NF-KB Targets
Relative mRNA Expression
NFKBIE
RELB
j
ATF4 Targets
Relative mRNA Expression
HOXB9
MAP3K8

**Figure S8:**

**Figure S8: (cont.)**

**a** Representative immunoblot showing the expression of the constructs used in the rescue experiments described in Figure 6. **b-g** Quantification of the immunoblot in Figure 6f. **h-j** qRT-PCR data shown in Figure 6h separated by pathway. Bars represent the average of at least three replicates, error bars are +/- SD. * $p < 0.05$, ** $p < 0.01$, *** $p < 0.001$. P-values determined by Dunnett’s test.

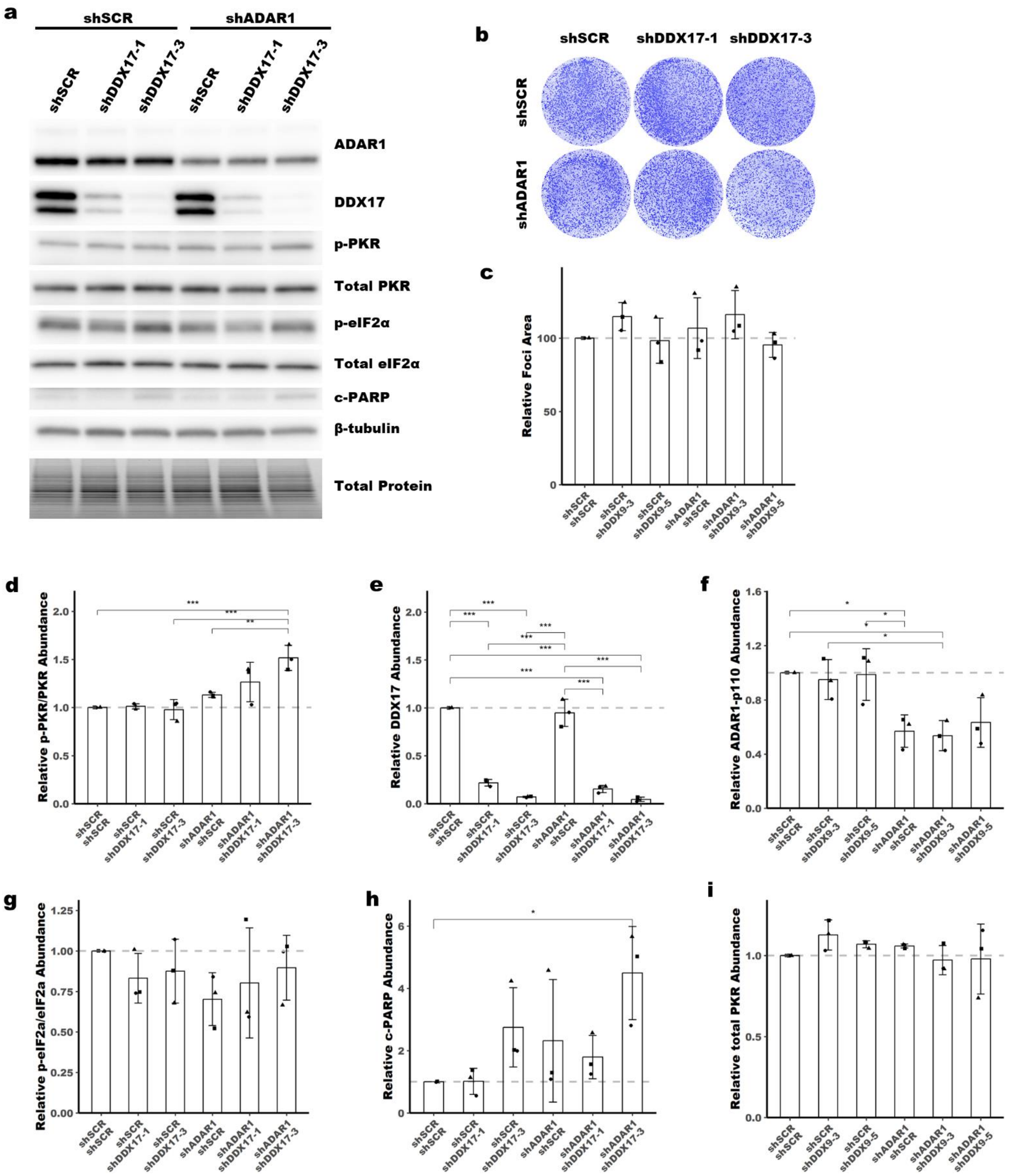

**Figure S9:**

**a** Representative immunoblot showing the phenotype of ADAR1 and/or DDX17 knockdown in SK-BR-3. Uncropped blots can be found in Source Data Figures. Protein abundance from the immunoblot in **a** was normalized by total protein abundance by quantification of the Stain Free gel image (bottom of panel). **b** Representative foci formation phenotype of ADAR1 and/or DD17 knockdown in SK-BR-3, quantification of relative foci area is shown in **c**. **d**-**i** Quantification of the immunoblot in **a**. Bars represent the average of at least three replicates, error bars are +/- SD. * p <0.05, ** p <0.01, *** p < 0.001.

**Figure S9: (cont.)**
P-values determined by one-way ANOVA with post-hoc Tukey. Comparisons between the two different shRNAs targeting DDX17 (shDDX17-1 and shDDX17-3) were not included for clarity.

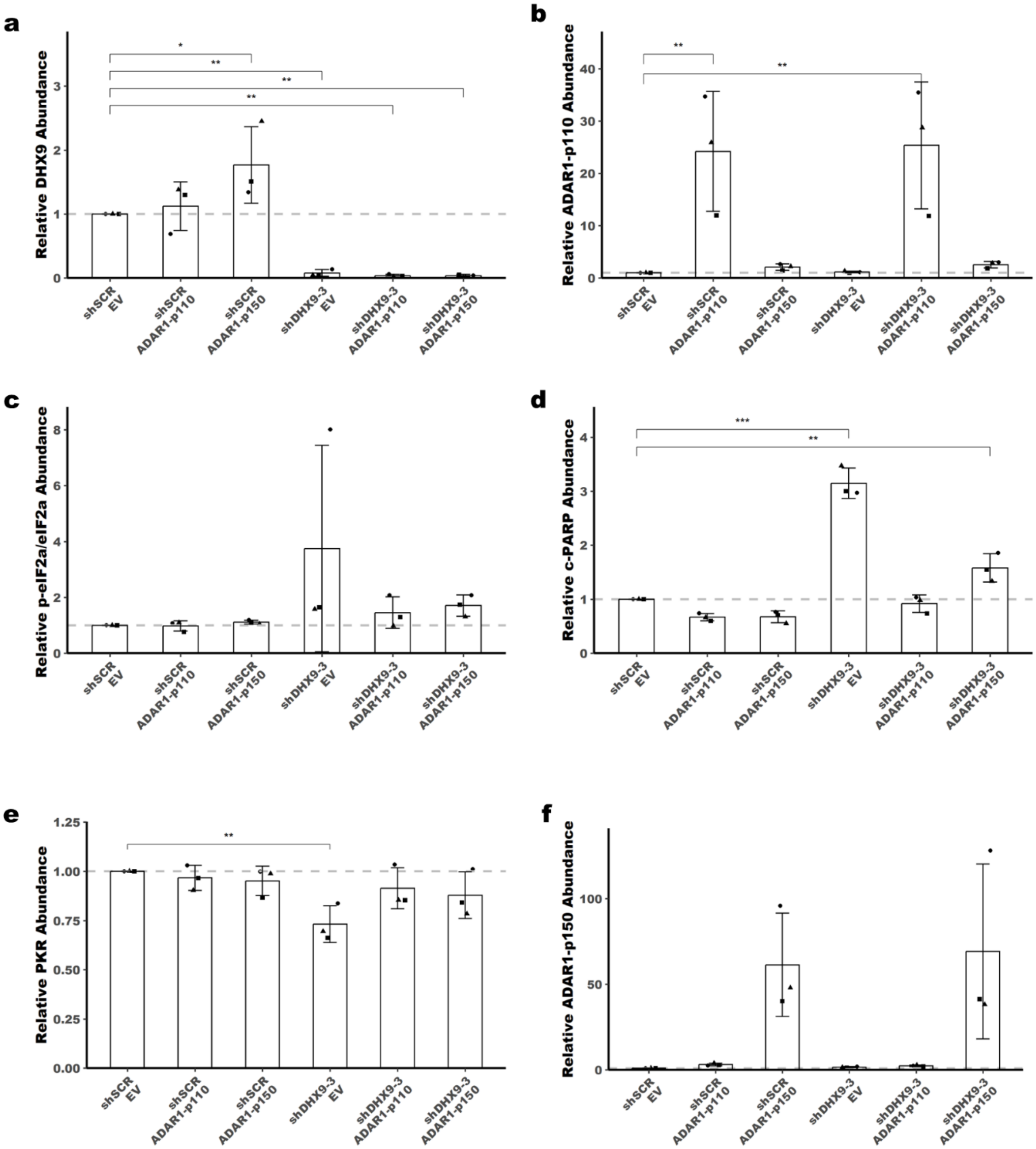


**Figure S10:**

**a**-**f** Quantification of the immunoblot in Figure 7a. Bars represent the average of at least three replicates, error bars are +/- SD. * $p < 0.05$, ** $p < 0.01$, *** $p < 0.001$. P-values determined by Dunnett’s test.

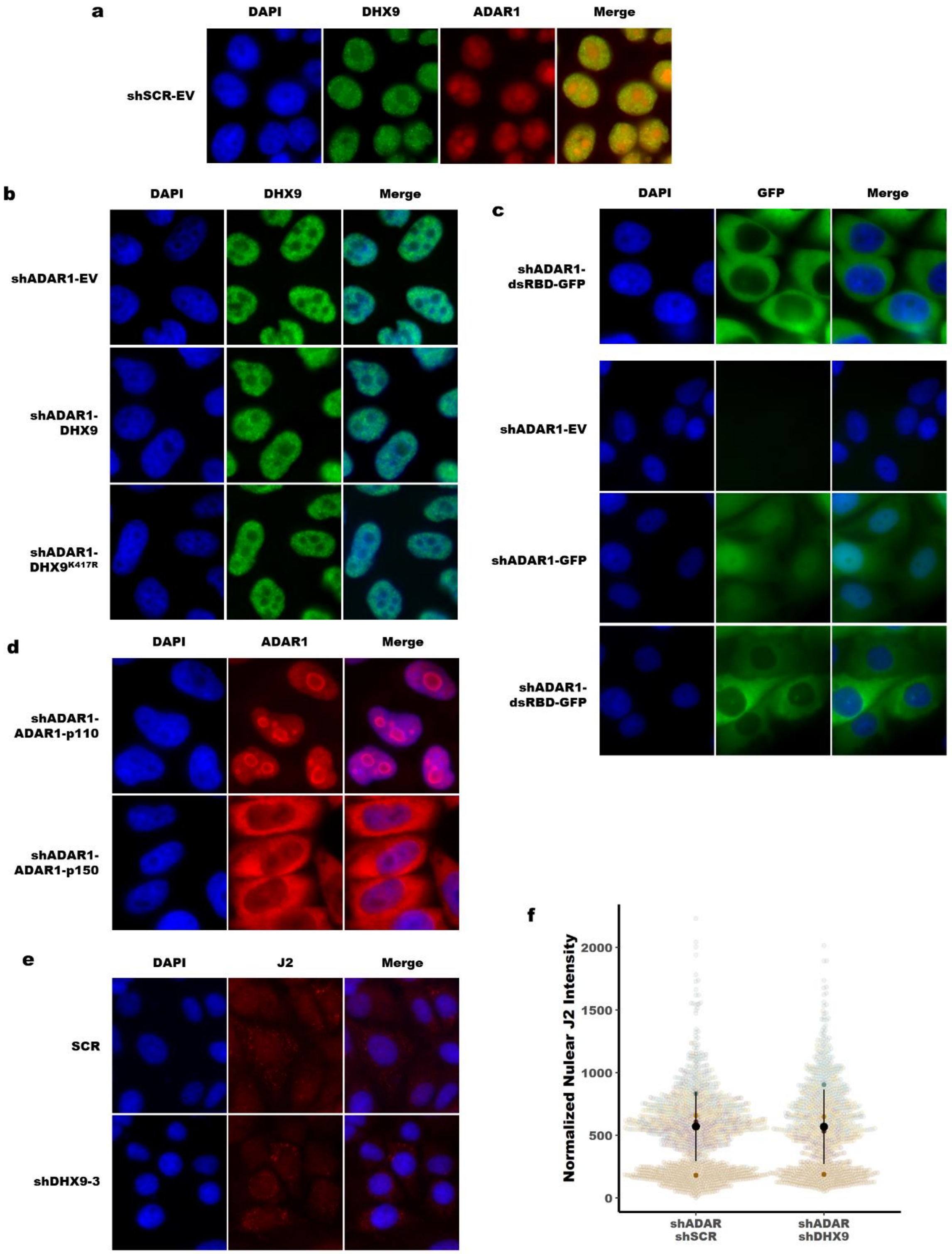


**Figure S11:**

Immunofluorescence for ADAR1 and DHX9 **a**, DHX9 **b**, GFP **c**, and ADAR1 **d** in SK-BR-3 infected with knockdown and overexpression constructs used in Figure 6. **e** Representative immunofluorescence for dsRNA with the J2 antibody following knockdown of ADAR1 (both conditions) and DHX9 (shDHX9-3). Nuclear J2 intensity is quantified for four replicates in panel **f**. The transparent colored dots (background) represent individual nuclei across three separate fields for each replicate. The opaque colored dots are the average for each replicate, colored by replicate to match the transparent dots. The black dot and lines represent the mean +/- SD.

Below are the source data figures.

Figure 1

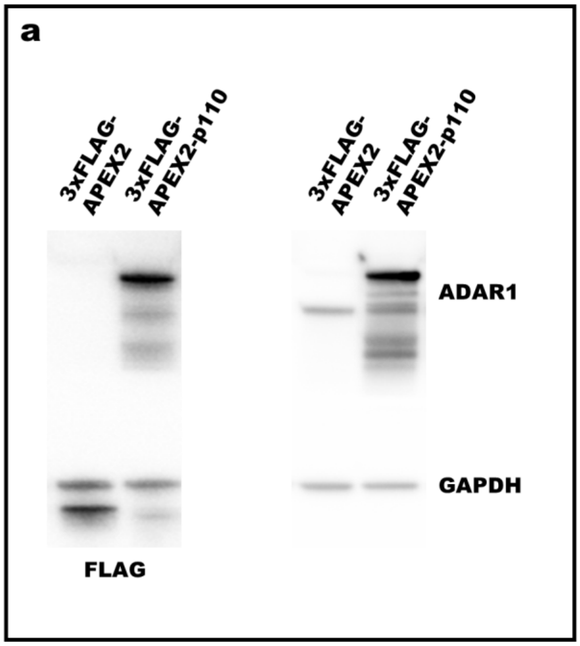

a
3xFLAG-APEX2
3xFLAG-APEX2-p110
3xFLAG-APEX2
3xFLAG-APEX2-p110
ADAR1
GAPDH
FLAG

Figure 2

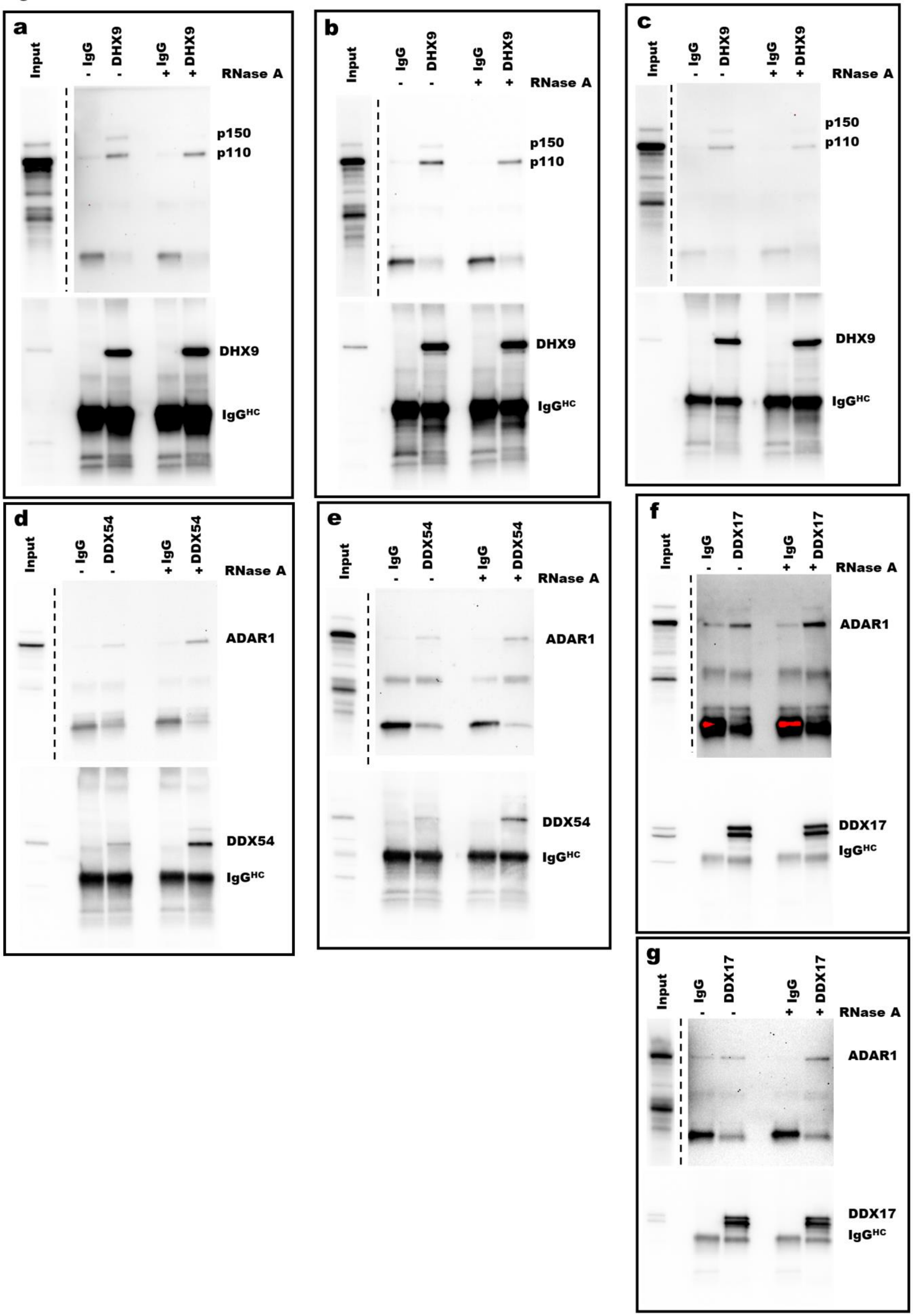
a
Input
- IgG
- DHX9
+ IgG
+ DHX9
RNase A
p150
p110
DHX9
IgG$^{HC}$
b
Input
IgG
DHX9
IgG
DHX9
- - + +
RNase A
p150
p110
DHX9
IgG$^{HC}$
c
Input
- IgG
- DHX9
+ IgG
+ DHX9
RNase A
p150
p110
DHX9
IgG$^{HC}$
d
Input
- IgG
- DDX54
+ IgG
+ DDX54
RNase A
ADAR1
DDX54
IgG$^{HC}$
e
Input
IgG
DDX54
IgG
DDX54
- - + +
RNase A
ADAR1
DDX54
IgG$^{HC}$
f
Input
IgG
DDX17
IgG
DDX17
- - + +
RNase A
ADAR1
DDX17
IgG$^{HC}$
g
Input
IgG
DDX17
IgG
DDX17
- - + +
RNase A
ADAR1
DDX17
IgG$^{HC}$

Figure 3

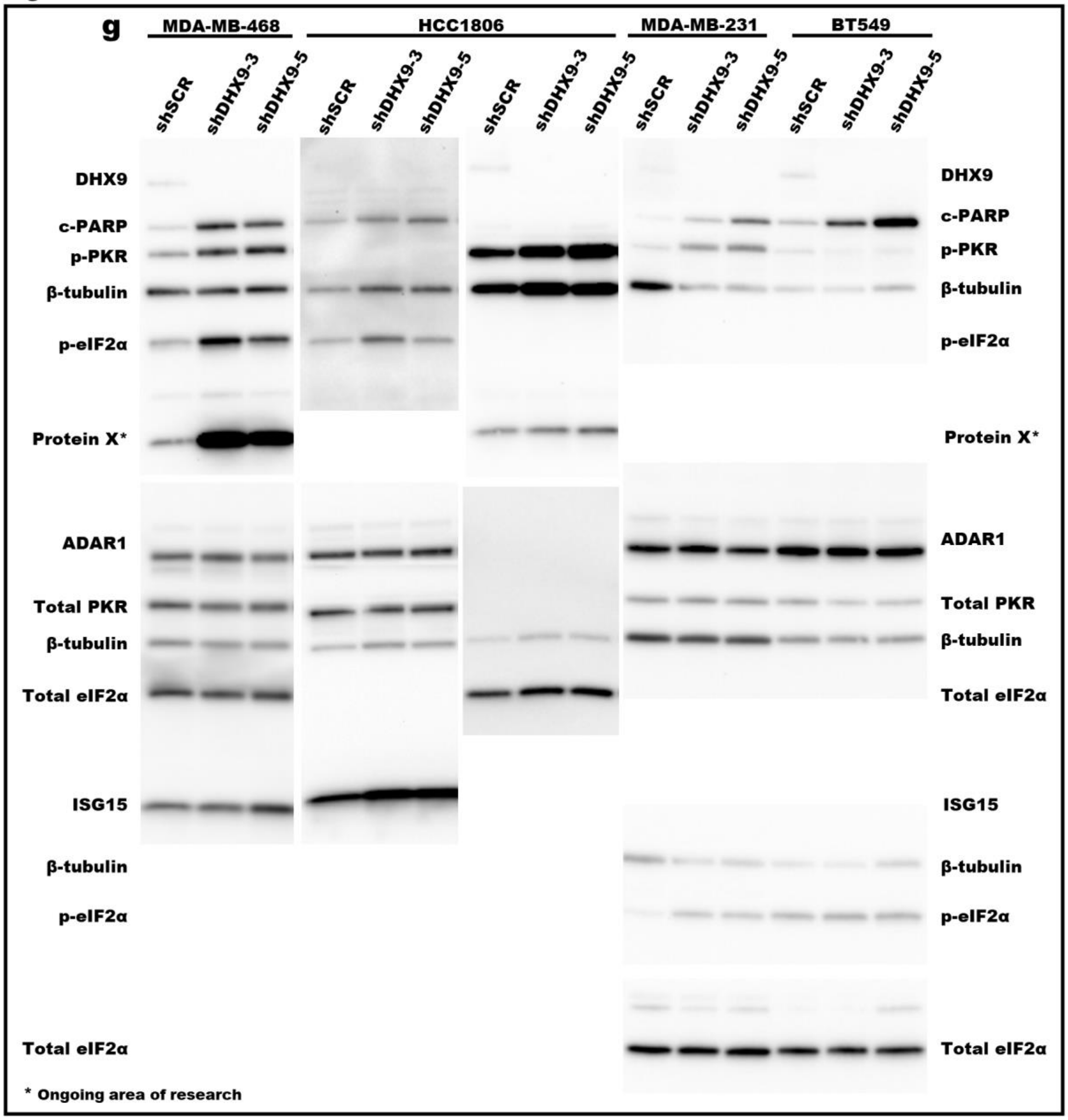


Extended Data Figure 3

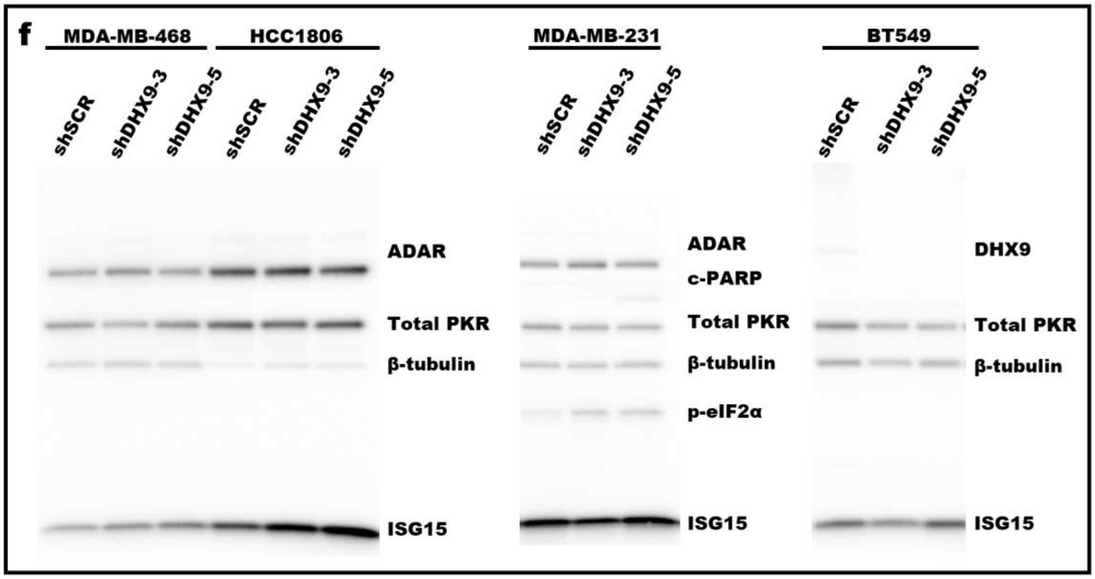

Figure 4

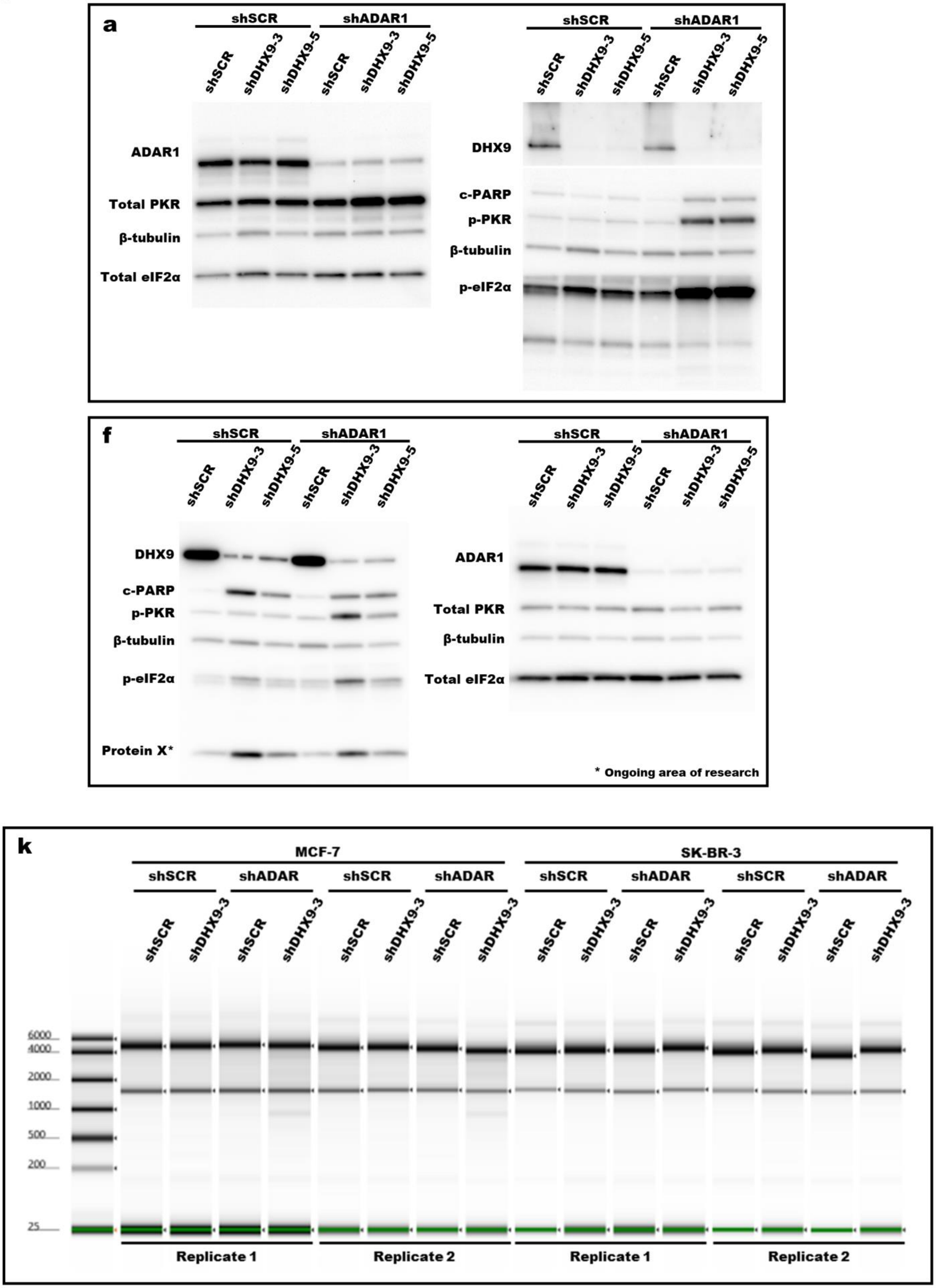
a
shSCR
shADAR1
shSCR
shDHX9-3
shDHX9-5
ADAR1
Total PKR
β-tubulin
Total eIF2α
DHX9
c-PARP
p-PKR
p-eIF2α
f
Protein X*
* Ongoing area of research
k
MCF-7
SK-BR-3
shADAR
6000
4000
2000
1000
500
200
25
Replicate 1
Replicate 2

Figure 4 and Extended Data Figure 4

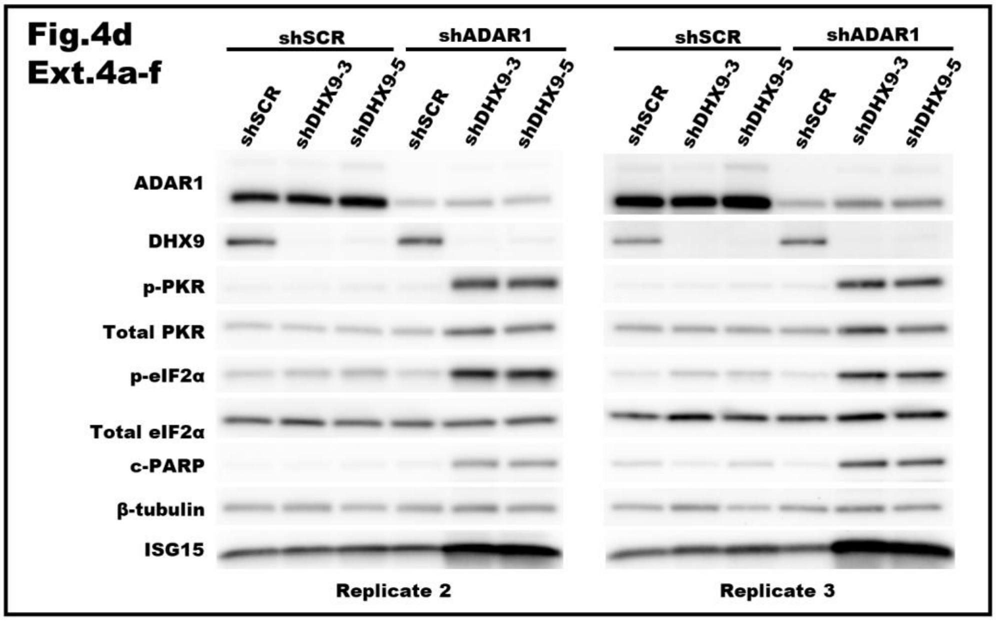


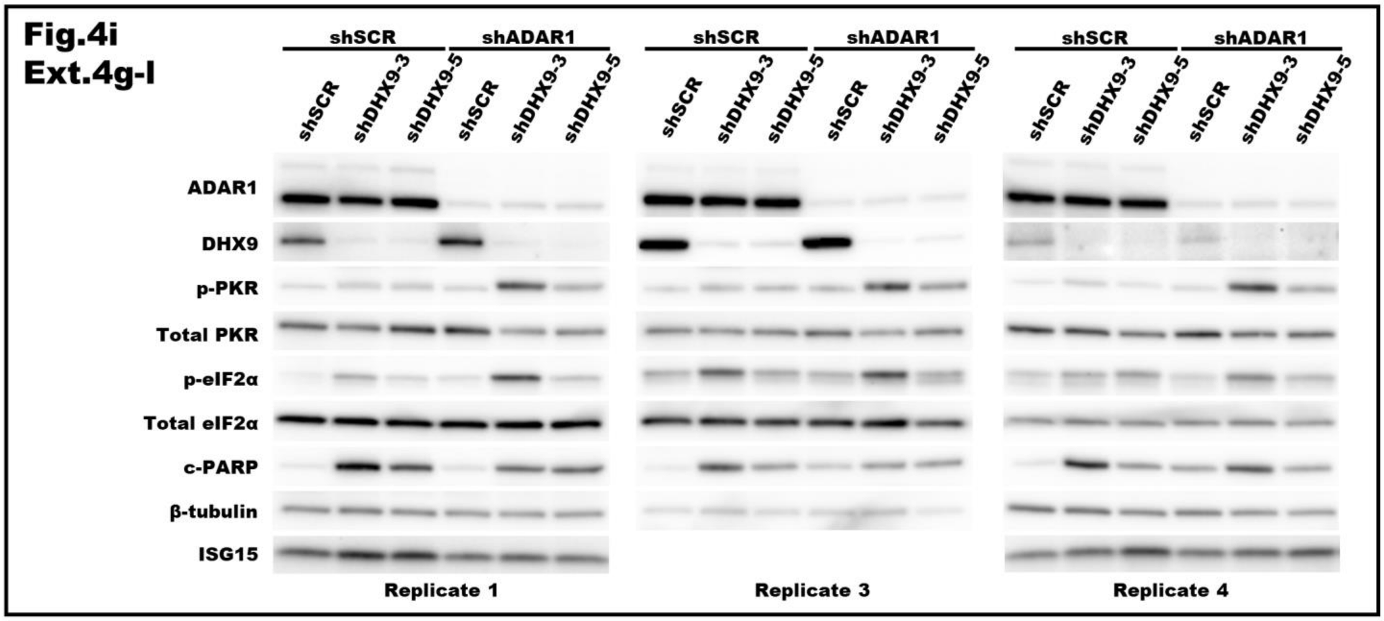

Figure 6 and Extended Data Figure 8

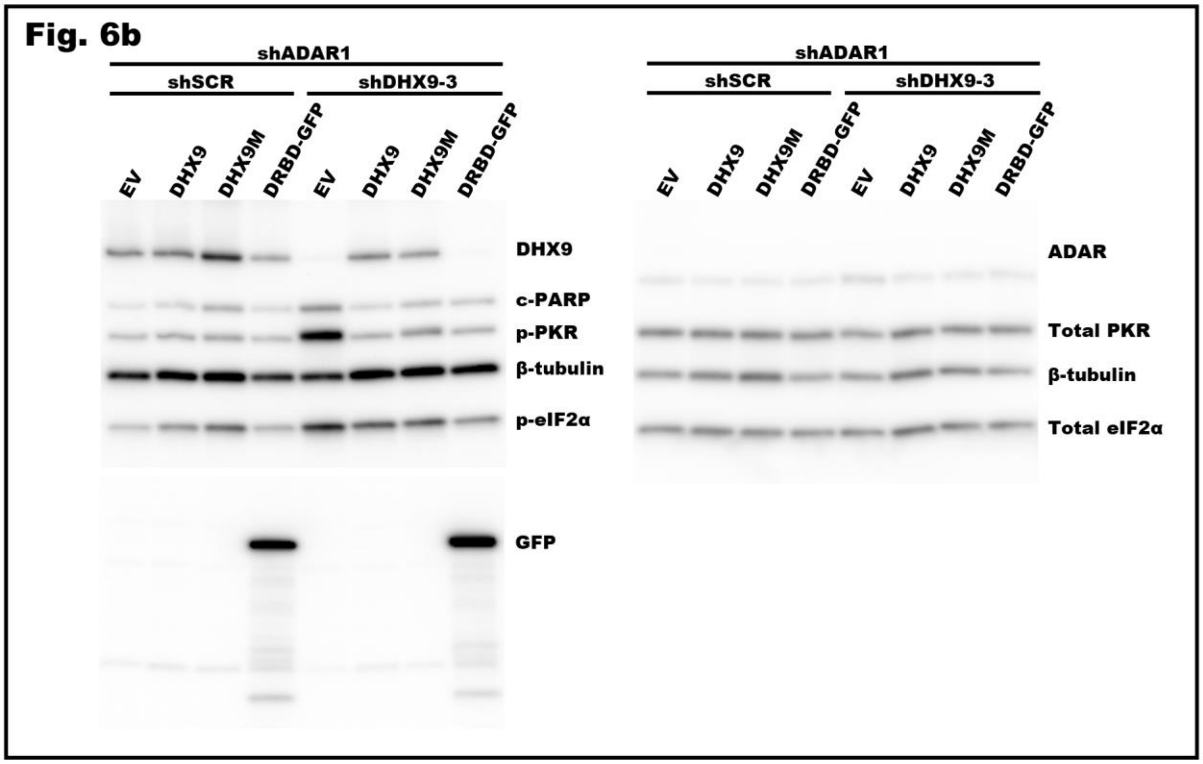


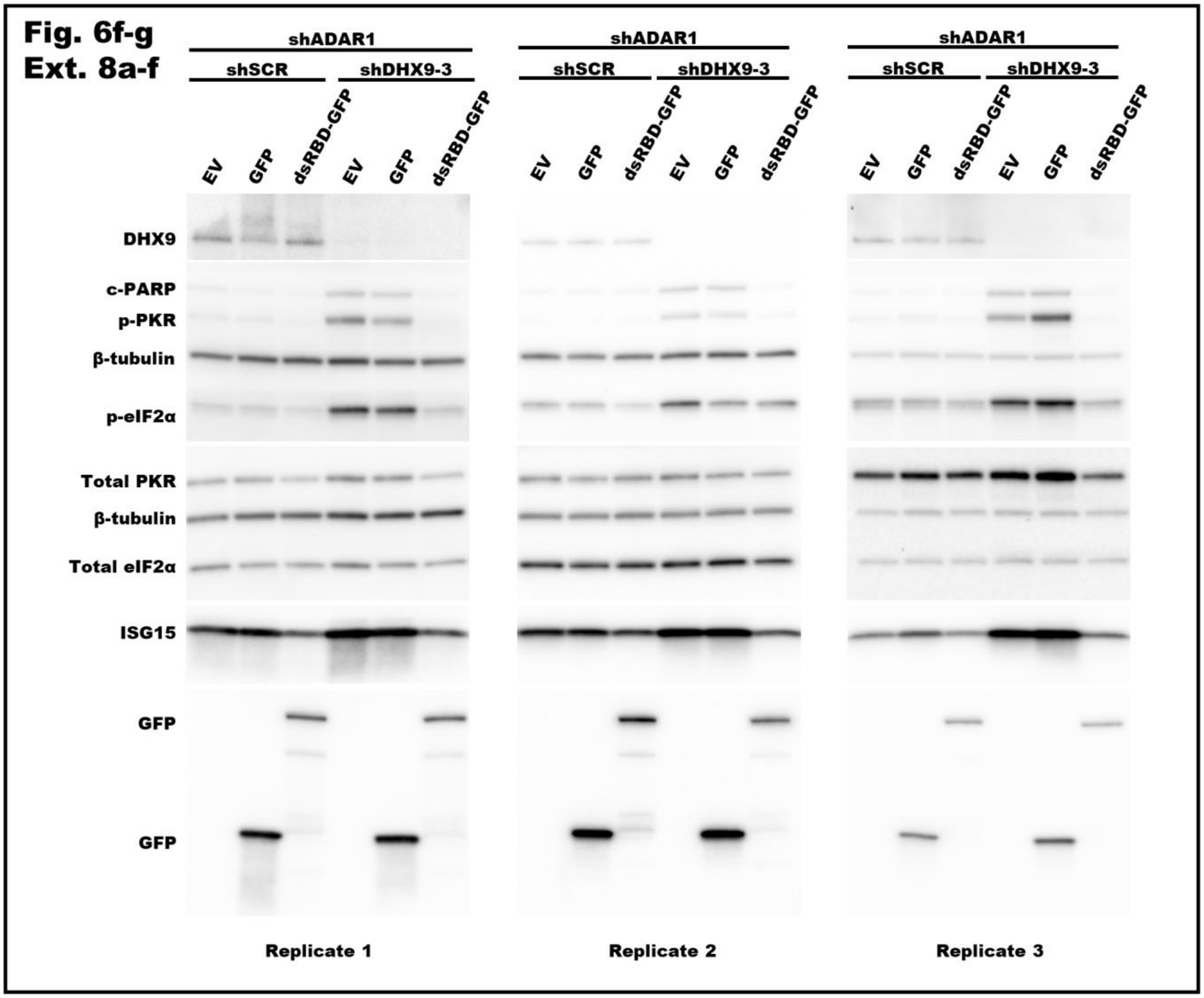

Figure 6 and Extended Data Figure 7

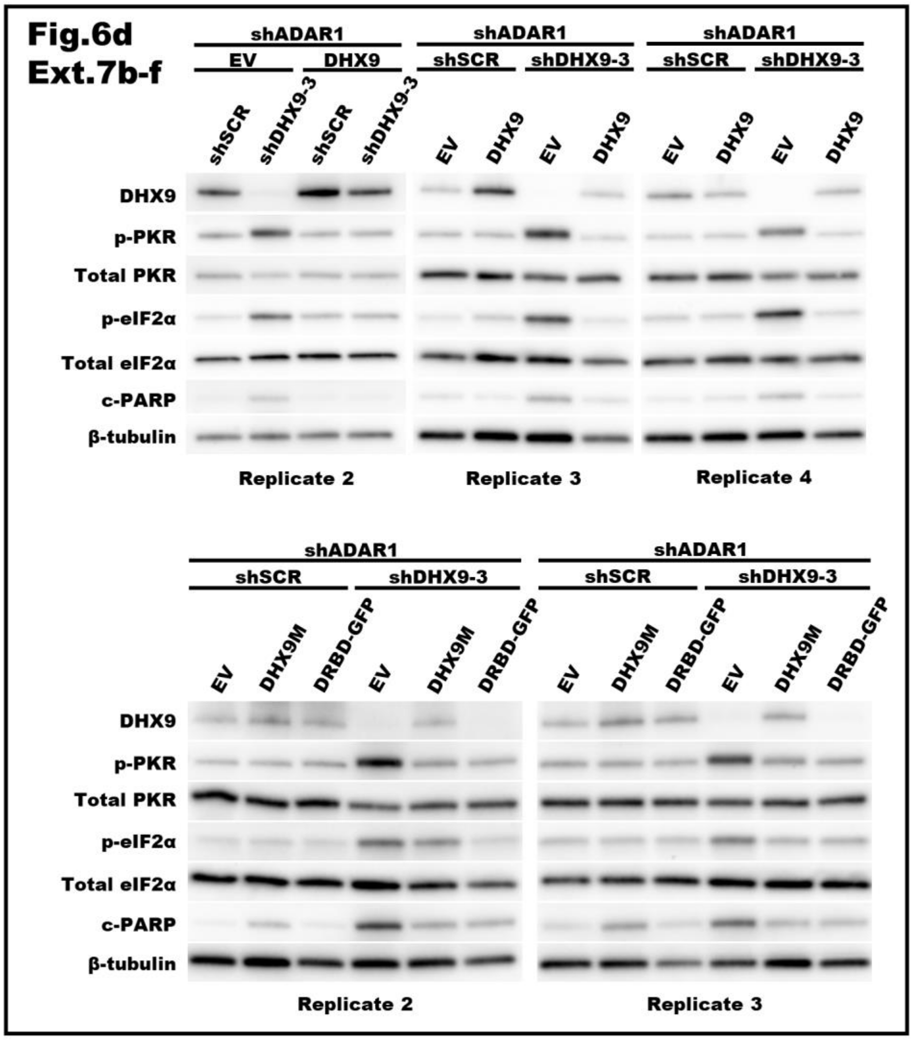

Figure 7 and Extended Data Figure 9

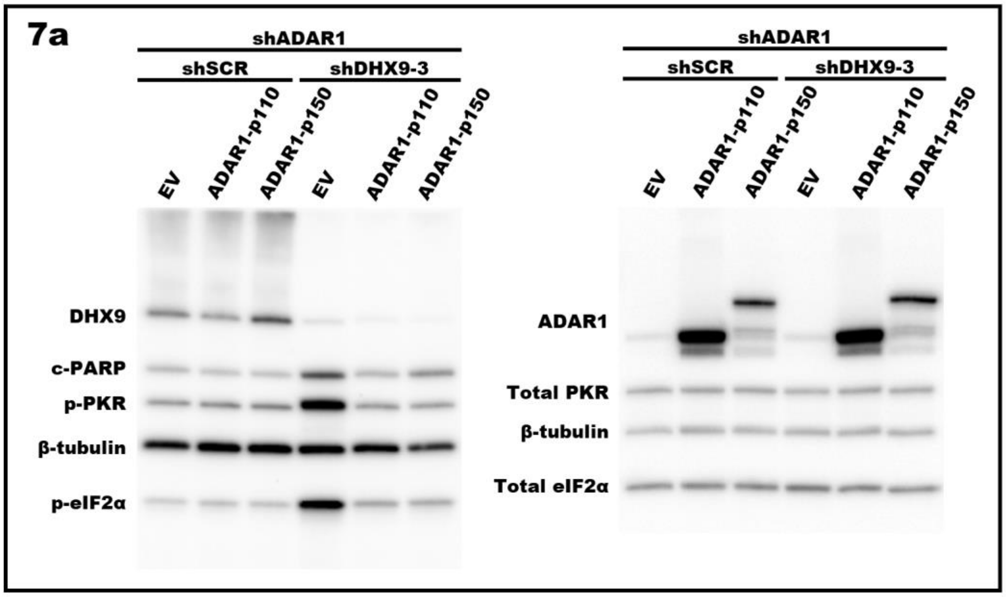


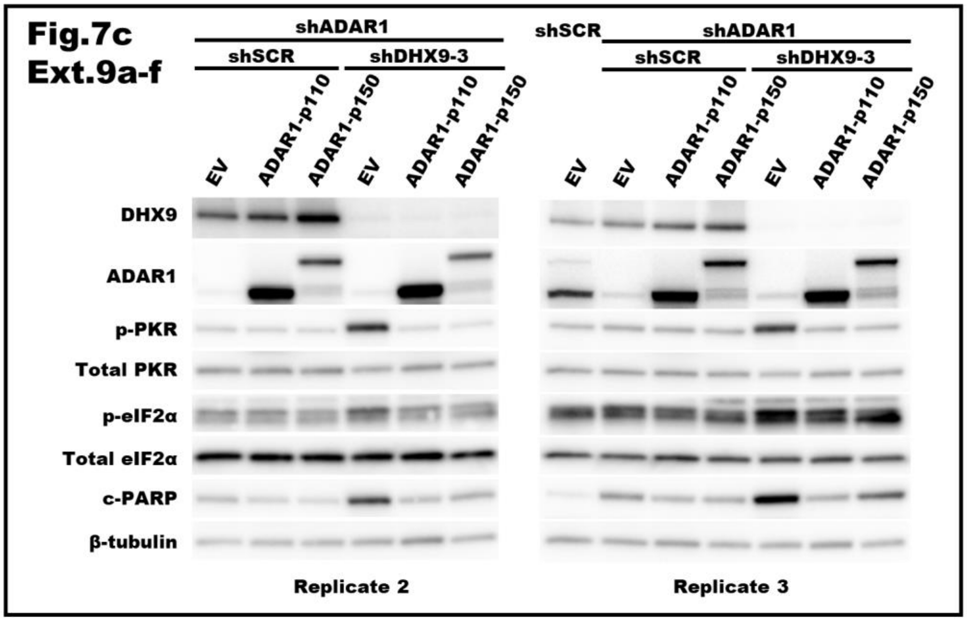

Extended Data Figure 10

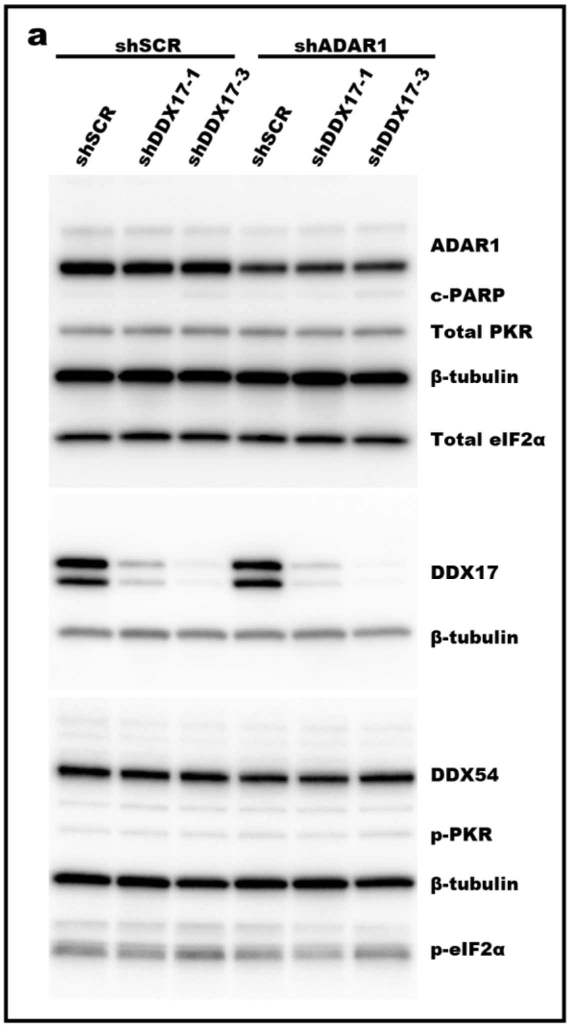
